# Structural basis of CHMP2A-CHMP3 ESCRT-III polymer assembly and membrane cleavage

**DOI:** 10.1101/2022.04.12.487901

**Authors:** Kimi Azad, Delphine Guilligay, Cecile Boscheron, Sourav Maity, Nicola De Franceschi, Guidenn Sulbaran, Gregory Effantin, Haiyan Wang, Jean-Philippe Kleman, Patricia Bassereau, Guy Schoehn, Wouter H Roos, Ambroise Desfosses, Winfried Weissenhorn

## Abstract

The endosomal sorting complex required for transport (ESCRT) is a highly conserved protein machinery that drives a divers set of physiological and pathological membrane remodeling processes. However, the structural basis of ESCRT-III polymers stabilizing, constricting and cleaving negatively curved membranes is yet unknown. Here we present cryo electron microscopy structures of membrane-coated CHMP2A-CHMP3 filaments of two different diameters at 3.3 and 3.6 Å resolution. The structures reveal helical filaments assembled by CHMP2A-CHMP3 heterodimers in the open ESCRT-III conformation, which generates a partially positive charged membrane interaction surface, positions short N-terminal motifs for membrane interaction and the C-terminal VPS4 target sequence towards the tube interior. Inter-filament interactions are electrostatic, which facilitate filament sliding upon VPS4-mediated polymer remodeling. Fluorescence microscopy as well as high speed atomic force microscopy imaging corroborate that CHMP2A-CHMP3 polymers and VPS4 can constrict and cleave narrow membrane tubes, thus acting as a minimal membrane fission machinery.

## Introduction

The endosomal sorting complex required for transport (ESCRT) machinery catalyzes many divergent membrane remodeling processes including the formation of multivesicular endosomes, cytokinesis, nuclear envelope reformation, membrane repair, autophagy, exosome biogenesis, neuronal pruning, dendritic spine maintenance, enveloped virus budding, release of peroxisomes and of recycling endosomes (Allison et al., 2013; Henne et al., 2013; Loncle et al., 2015; Mast et al., 2018; Olmos and Carlton, 2016; Sadoul et al., 2018; Scourfield and Martin-Serrano, 2017; Votteler and Sundquist, 2013; Zhen et al., 2021).

Common to all ESCRT-catalyzed processes in eukaryotes, archaea and bacteria is the recruitment of ESCRT-III proteins that polymerize to generate and/or to stabilize membranes with either flat, negatively or positively curved geometries (Bertin et al., 2020; Caillat et al., 2019; Gupta et al., 2021; Junglas et al., 2021; Liu et al., 2021; McCullough et al., 2018; Moser von Filseck et al., 2020; Pfitzner et al., 2021). The principal function of the polymers is to induce membrane constriction via outside-in fission of tubular structures with ESCRT-III protein coats on the outside of a membrane tube or inside-out fission with ESCRT-III polymers assembled within membrane neck/tube structures formed during vesicle and virus budding or at the cytokinetic midbody (Caillat et al., 2019; Harker-Kirschneck et al., 2022; McCullough et al., 2018; Nguyen et al., 2020; Pfitzner et al., 2021; Remec Pavlin and Hurley, 2020).

Humans express eight ESCRT-III proteins that can comprise several isoforms per member, corresponding to seven homologues in *S. cerevisiae* (in parentheses) named CHMP1A/B (Did2), CHMP2A/B (Vps2), CHMP3 (Vps24), CHMP4A/B/C (Snf7), CHMP5 (Vps60), CHMP6 (Vps20), CHMP7 and CHMP8/IST1 (Ist1) (McCullough et al., 2018). *S. cerevisiae* assembles two ESCRT-III subcomplexes, Vps20/Snf7 that in turn recruits Vps24/Vps2 (Babst et al., 2002) consistent with CHMP4 recruiting CHMP3 and CHMP2A (Morita et al., 2011). Notably, Vps24 (CHMP3) and Vps2 (CHMP2) have been suggested to block Snf7 (CHMP4) polymerization and cap ESCRT-III assembly prior to recycling (Saksena et al., 2009; Teis et al., 2008). Consistent with the idea of a core ESCRT-III, HIV-1 budding can be catalyzed with a minimal set of one CHMP4 and one CHMP2 isoform (Morita et al., 2011). Although CHMP3 is not strictly required, truncated versions thereof exert a potent dominant negative effect on HIV-1 budding (Zamborlini et al., 2006) and CHMP3 synergizes HIV-1 budding efficiency with CHMP2A but not with CHMP2B (Effantin et al., 2013). Thus ESCRT-III CHMP4, CHMP2 and CHMP3 constitute a minimal machinery that together with VPS4 catalyzes membrane fission from within membrane necks as suggested by *in vitro* reconstitution (Schoneberg et al., 2018).

ESCRT-III proteins adopt a closed conformation in the cytosol (Bajorek et al., 2009; Muziol et al., 2006; Xiao et al., 2009). Membrane recruitment via ESCRT-I, ESCRT-II or Alix/Bro1 (Im et al., 2009; McCullough et al., 2008; Pineda-Molina et al., 2006; Tang et al., 2016) is thought to induce ESCRT-III activation, which entails opening of the closed conformation (Lata et al., 2008a; Shim et al., 2007; Zamborlini et al., 2006) to an extended open polymerization-competent conformation as first shown for CHMP1B (McCullough et al., 2015). The CHMP1B polymer stabilizes positively curved membranes and can co-polymerize with IST1 in the closed conformation thereby forming an outer layer on top of the inner open conformation CHMP1B layer (McCullough et al., 2015), whose interplay leads to membrane tube thinning and cleavage (Cada et al., 2022; Nguyen et al., 2020).

CHMP4 homologues, Snf7 and shrub, adopt similar open conformations within crystalline polymers (McMillan et al., 2016; Tang et al., 2015). Notably *in vitro* CHMP4 polymers interact with flat membranes, (Chiaruttini et al., 2015; Mierzwa et al., 2017; Pires et al., 2009) and stabilize positively curved membranes in the presence of CHMP2A (Vps2) and CHMP3 (Vps24) (Bertin et al., 2020; Moser von Filseck et al., 2020). Furthermore, CHMP4 was proposed to interact with negatively curved membranes (Lee et al., 2015) and CHMP4 spirals have been imaged within membrane tubes *in vivo* (Cashikar et al., 2014; Hanson et al., 2008) leading to the model of ESCRT-III spiral springs assembling on flat membranes driving membrane deformation (Chiaruttini et al., 2015).

The conservation of the structural principle of the open ESCRT-III conformation is further underlined by the structures of plastid and bacterial membrane repair proteins Vipp1 and PspA, which both stabilize positively curved membranes and corroborate the conservation of the ESCRT-III machinery throughout all kingdoms of life (Gupta et al., 2021; Junglas et al., 2021; Liu et al., 2021).

While these structures demonstrate filament formation to stabilize positively curved membranes only low resolution models of filaments stabilizing negatively curved membranes have yet been imaged revealing single and multi-stranded polymers *in vitro* (Bajorek et al., 2009; Bertin et al., 2020; Chiaruttini et al., 2015; Dobro et al., 2013; Henne et al., 2012; Lata et al., 2008b; Mierzwa et al., 2017; Moriscot et al., 2011; Moser von Filseck et al., 2020; Pires et al., 2009) and *in vivo* (Bodon et al., 2011; Cashikar et al., 2014; Goliand et al., 2018; Guizetti et al., 2011; Hanson et al., 2008; Mierzwa et al., 2017; Sherman et al., 2016). Consistent with their central role in membrane fission catalyzed from within membrane necks, ESCRT-III CHMP2A and CHMP3 form helical tubular structures with defined diameters *in vitro,* which have been suggested to stabilize negative membrane curvature (Effantin et al., 2013; Lata et al., 2008b). VPS4 constricts these filaments producing dome-like end caps prior to complete polymer disassembly *in vitro* (Maity et al., 2019) in agreement with permanent ESCRT-III turn-over *in vivo* (Adell et al., 2014; Adell et al., 2017; Mierzwa et al., 2017). Notably, *S. cerevisiae* Vps2 and Vps24 form similar helical tubes that, however, seem to require Snf7 for polymerization (Henne et al., 2012).

Because most ESCRT-catalyzed processes act on negatively curved membranes to catalyze inside-out membrane fission, we set out to determine the structural basis of ESCRT-III stabilizing negatively curved membranes. We reconstituted CHMP2A-CHMP3 polymers within membrane tubes, solved their structure by cryo electron microscopy and show that VPS4 can indeed constrict CHMP2A-CHMP3 membrane tubes to the point of fission, corroborating that CHMP2A and CHMP3 form a minimal membrane fission complex powered by VPS4 and ATP.

## Results

### Structure of the CHMP2A-CHMP3 polymer assembled within membrane tubes

CHMP3 full length (residues 1-222) and C-terminally truncated CHMP2A (residues 1-161) were assembled into helical tubular structures as described (Lata et al., 2008b). After removal of the N-terminal tags, the tubular structures were coated with lipid bilayer, which tightly associated with the protein layer as shown by cryo electron microscopy (cryo-EM) (**Figure S1A**). 2D classification of the manually picked tube-like structures generated a dataset composed of 5 different diameters ranging from 380 Å to 490 Å, with the 410 Å (51.4%) and 430 Å (31.2%) diameters representing the most populated classes (**Figure S1B**). The power spectra of the segments of the class averages of the two diameters (**Figures S1C and D**) were then employed to explore possible helical symmetries and combined with helical real-space reconstruction (He and Scheres, 2017) to validate the symmetry parameters. This revealed that the 410 Å and 430 Å diameter tubes are formed by elementary helices composed of respectively 6.3 and 6.6 units per turn with a small pitch of 9 Å for the 410 Å diameter and 18 Å for the 430 Å diameter. The latter displays an additional C2 symmetry around the helical axis, explaining the doubling of the pitch (**Figures S1C and D**). Although, the asymmetric units along the elementary helix are not biochemically connected, they translate into filaments with extended interaction surfaces between subsequent asymmetric units. The symmetry parameters of the filaments are relatively similar for both diameters (Rise/Twist of 8.227 Å /16.877° and 8.641 Å /17.701° for the 410 Å and 430 Å diameters, respectively), and therefore likely represent the preferred polymerization mode of the repeating unit, the CHMP2A-CHMP3 heterodimers (**Figures S1C and D**).

Six filaments (2*3 in the case of the C2 symmetric helix) form left-handed six-start helices with helical pitches of ∼175 Å for both diameters (**Figures 1A** **and** **B**). The 430 Å filament is composed of 21.33 units per turn (**Figures 1A** **and** **B**) and the 410 Å filament contains 20.33 units per turn (**Figures 1C** **and** **D**), indicating a helical repeat after three turns and similar inter-filament interactions along the helical axis. Comparison of the 430 and 410 Å demonstrates that removal of one heterodimer reduces the tube diameter by 20 Å.

**Figure 1:**
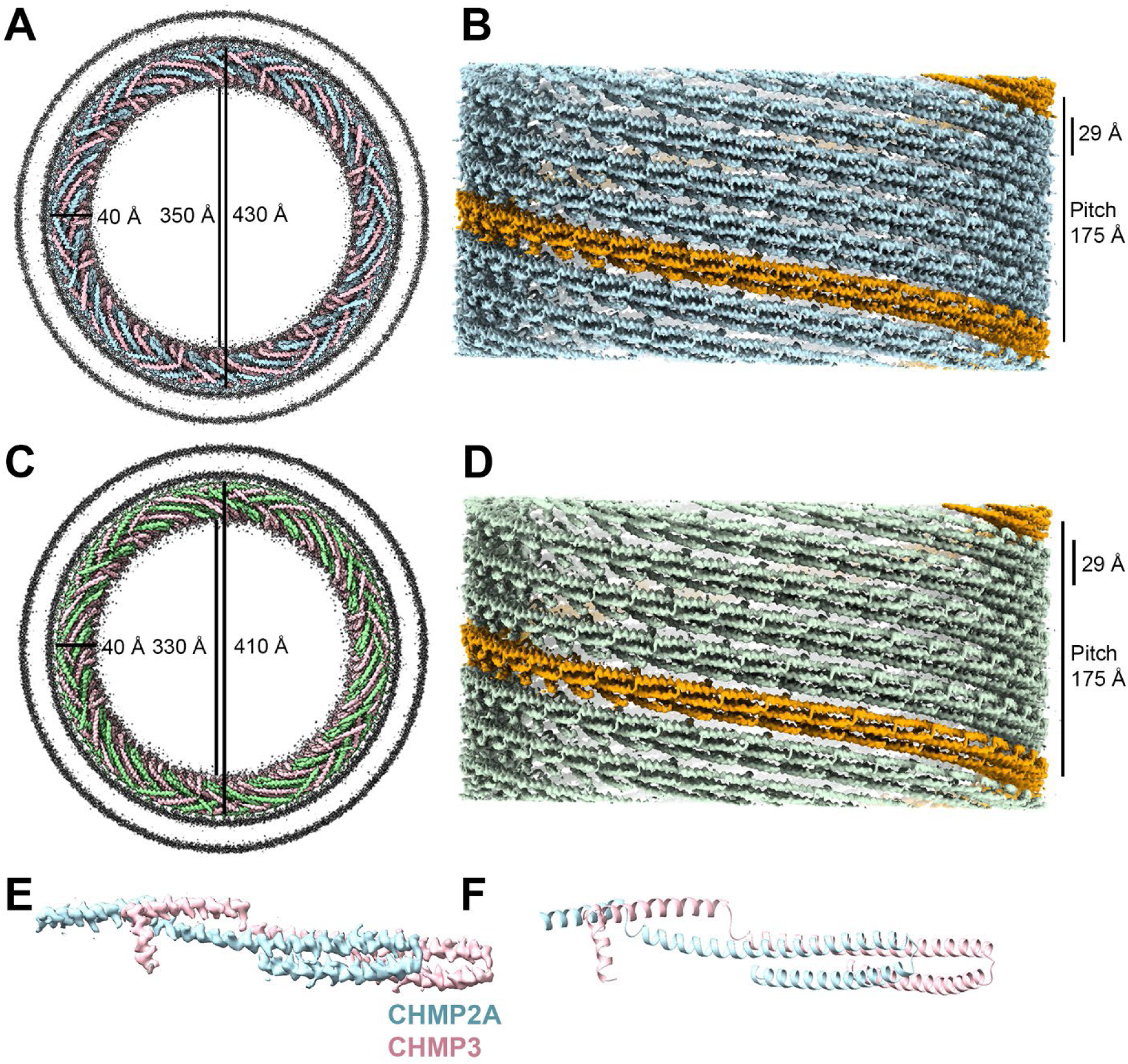
Cryo-EM structure of CHMP2A-CHMP3 membrane-coated helical polymers. **(A)** Density map of the reconstructed 430 Å diameter CHMP2A-CHMP3 membrane tube with the top view looking down the helical axis. The helical arrangement of CHMP2A (light blue) and CHMP3 (pink) inside the bilayer membrane (dark grey) is shown. The thickness, and the inner and outer diameter of the helical protein tube are also marked in Å. **(B)** Side view of the helical polymer without the lipid membrane. One left-handed filament is indicated in orange, and the thickness of one filament and the pitch of the helical assembly are also marked. **(C)** Density map of the reconstructed 410 Å diameter CHMP2A-CHMP3 membrane tube with the top view looking down the helical axis. The helical arrangement of CHMP2A (green) and CHMP3 (pink) inside the bilayer membrane (dark grey) is shown. The thickness, and the inner and outer diameter of the helical protein tube are indicated in Å. **(D)** Side view of the helical polymer without the lipid membrane. One left-handed filament is indicated in orange, and the thickness of one filament and the pitch of the helical assembly are indicated. **(E)** Cryo-EM density of the single repeating unit of the 430 Å diameter polymer formed by the heterodimer of CHMP2A (light blue) and CHMP3 (pink) is indicated. **(F)** Ribbon representation of the atomic model of CHMP2A (light blue) - CHMP3 (pink) heterodimer.

The three-dimensional (3D) helical reconstruction shows an overall resolution of 3.6 Å for the 410 Å and 3.3 Å for the 430 Å diameter (**Figure 1; Figure S2 and Figure S3A**) with local resolutions ranging from 3.3 Å to 4.6 Å for the 430 Å diameter and 3.6 to 5.2 Å for the 410 Å diameter (**Figure S3B and C).** The map of the 430 Å diameter was employed (**Figure 1E**) to build the atomic model of the repeating unit of the filament, formed by the CHMP2A-CHMP3 heterodimer revealing both protomers in the open ESCRT-III conformation (**Figure 1F** **and Figure S3D, Table S1**).

Comparison of the closed (Bajorek et al., 2009; Muziol et al., 2006) and open CHMP3 conformations showed the conformational transitions upon CHMP3 activation, which involves extension of the helical hairpin (residues P12-A101) that is identical in both conformations (r.m.s.d. of 1.082 Å) by a linker and helix 3 (L117). The following short connection forms an elbow and translates helix 4 by ∼10 Å positioning helix 4 in a 140° angle with respect to the hairpin axis. Helix 4 is composed of the closed conformation helix 4 and most of the disordered linker connecting to helix 5 via a 90° kink at positions M151 to D152 (**Figure 2A**). The remaining CHMP3 residues 170 to 220 are flexible and disordered in the structure. Both CHMP3 and CHMP2A open conformations are similar as superposition of their Cα atoms revealed an r.m.s.d. of 0.934 Å (**Figure S4A**), suggesting that CHMP2A can fold into the same closed conformation structure as CHMP3.

**Figure 2:**
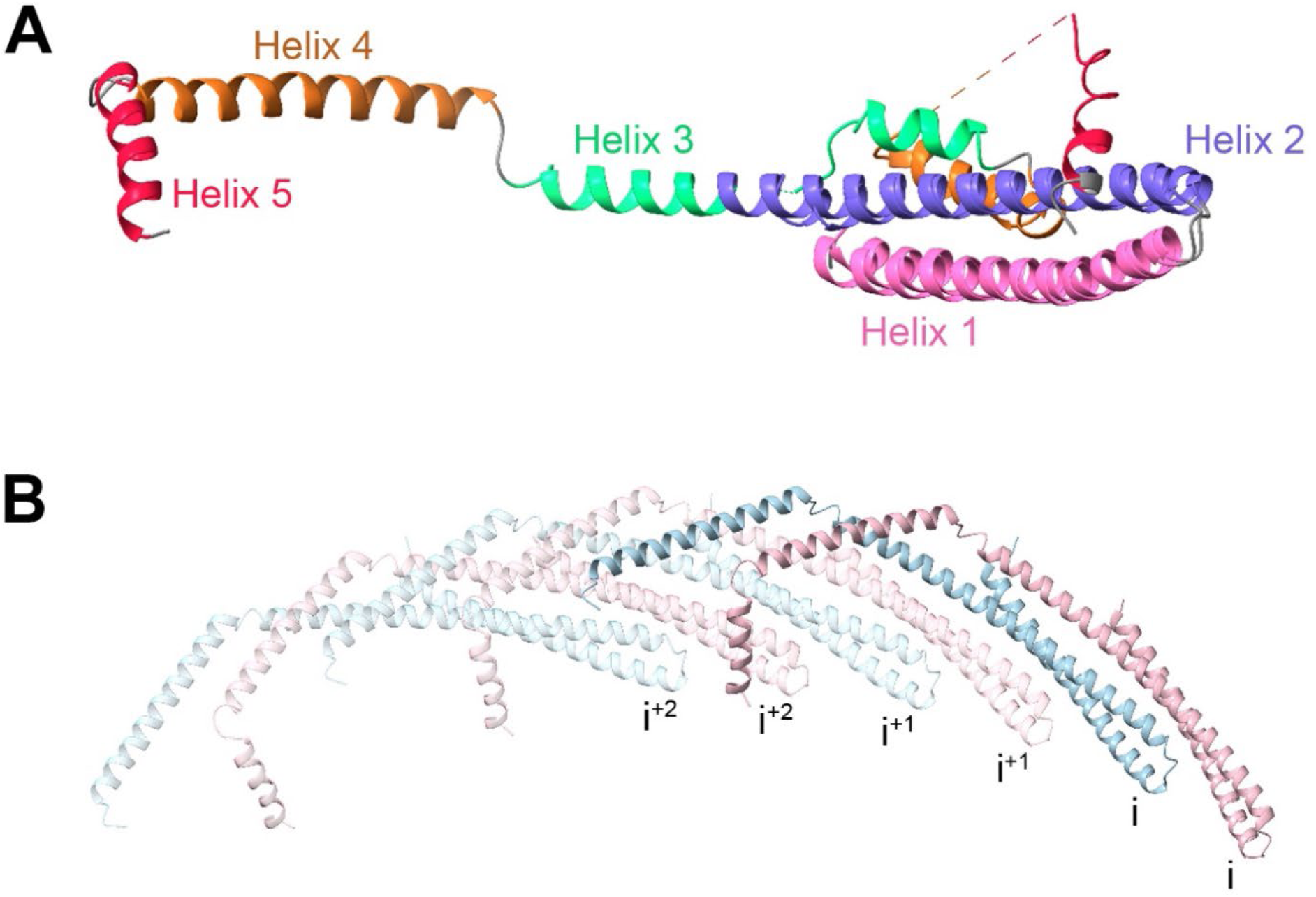
Atomic model and architecture of the CHMP2A-CHMP3 helical polymer. **(A)** Ribbon diagram of Cα superposition of the closed and open CHMP3 conformations (Helix 1: pink, Helix 2: purple, Helix 3: light green, Helix 4: brown, Helix 5: red). **(B)** Three interlocked copies of CHMP2A-CHMP3 heterodimer are shown as ribbons. Helix 4 and 5 of CHMP3 (pink) interact with four subsequent protomers. Helix 4 of CHMP2A (light blue) also makes similar interactions.

The repeating unit of the filament is the CHMP2A-CHMP3 heterodimer formed by parallel interaction of their hairpins with the CHMP2A hairpin tip shifted by six helical turns with respect to CHMP3 (**Figure 2B**). The heterodimer interaction covers 2026 Å^2^ of CHMP2A and 1997 Å^2^ of CHMP3 surfaces involving 55 and 51 interface residues, respectively. The structure of the heterodimer was further confirmed by mutagenesis. Introducing pairs of cysteine demonstrated that CHMP2A_D57C together with CHMP3_S75C and CHMP2A_N18C together with CHMP3_V110C (**Figure S5A**) assembled into disulfide-linked heterodimers upon polymerization into tube-like structures as shown by SDS-PAGE analyses and negative staining EM (**Figure S5B and C**). Furthermore, mutagenesis of CHMP2A-CHMP3 interface residues (**Figure S5D**) prevented polymerization as expected (**Figure S5E**). The principle of the heterodimer hairpin stacking is employed to assemble the filament, which is further stabilized by lateral interactions of CHMP3i elbow helix 4 with CHMP2Ai, CHMP3i^+1^ and CHMP2Ai^+1^. In addition, CHMP3i helix 5 interacts with the tip of the CHMP3i^+2^ hairpin (**Figure 2B**), as observed in the closed conformation (**Figure 2A**) (Bajorek et al., 2009; Muziol et al., 2006). Similar to CHMP3, CHMP2A helix 4 exerts the same domain exchange interactions (**Figure 2B**). Cα superposition of the open CHMP3 conformation revealed the closest match with the *S. cerevisiae* Snf7 protomer (**Figure S4B**) and considerable differences with CHMP1B, Vipp1, PspA and Vps24 (Gupta et al., 2021; Huber et al., 2020; Junglas et al., 2021; Liu et al., 2021; McCullough et al., 2015) (**Figure S4C-F**). Notably, different hairpin interactions and orientations of the helical arms upon polymerization determine the filament geometry that leads to positively curved membrane interaction by CHMP1B, Vipp1 and PspA (Gupta et al., 2021; Junglas et al., 2021; Liu et al., 2021; McCullough et al., 2015) underlining the extensive structural plasticity of ESCRT-III proteins.

### CHMP2A-CHMP3 polymer interaction with membrane

The CHMP2A-CHMP3 polymer is tightly associated with the lipid bilayer (**Figure S1E**) and both CHMP2A and CHMP3 expose the same regions to the membrane. The polymerization mode positions the N-terminal regions of both CHMP2A and CHMP3 at the membrane interface. Although CHMP3 residues 1-10 and CHMP2A residues 1-7 are disordered, they are both oriented by conserved prolines towards the lipid bilayer (**Figure 3A**) consistent with previous suggestions that short amphipathic N-terminal helices insert into the bilayer (Bodon et al., 2011; Buchkovich et al., 2013). The main membrane interaction surfaces locate to the elbow formed by helices 3 and 4 (residues K104 to R131) exposing six basic residues of CHMP2A (K104, K108, R115, K118, K124, R131) and five basic residues of CHMP3 (K106, K112, K119, K132, K136) prone to interact with negative charges of the membrane (**Figure 3B**). The electrostatic potential map shows in addition to the stretch of basic surfaces some negative and non-charged regions of the outer polymer surface (**Figure 3C**). Most of the basic residues are conserved in *S. cerevisiae* Vps2 and Vps24 (**Figure S6A**). Notably, alanine mutagenesis of some CHMP3 basic residues within the membrane interaction surface did not interfere with CHMP2A-CHMP3 polymerization *in vitro* (**Figure S7A**) nor did they affect the dominant negative effect of C-terminally truncated CHMP3 on VLP release (**Figure S7B**), indicating that membrane binding is complex and not only electrostatic, consistent with plasma membrane localization of the CHMP3 mutant (**Figure S7C**).

**Figure 3:**
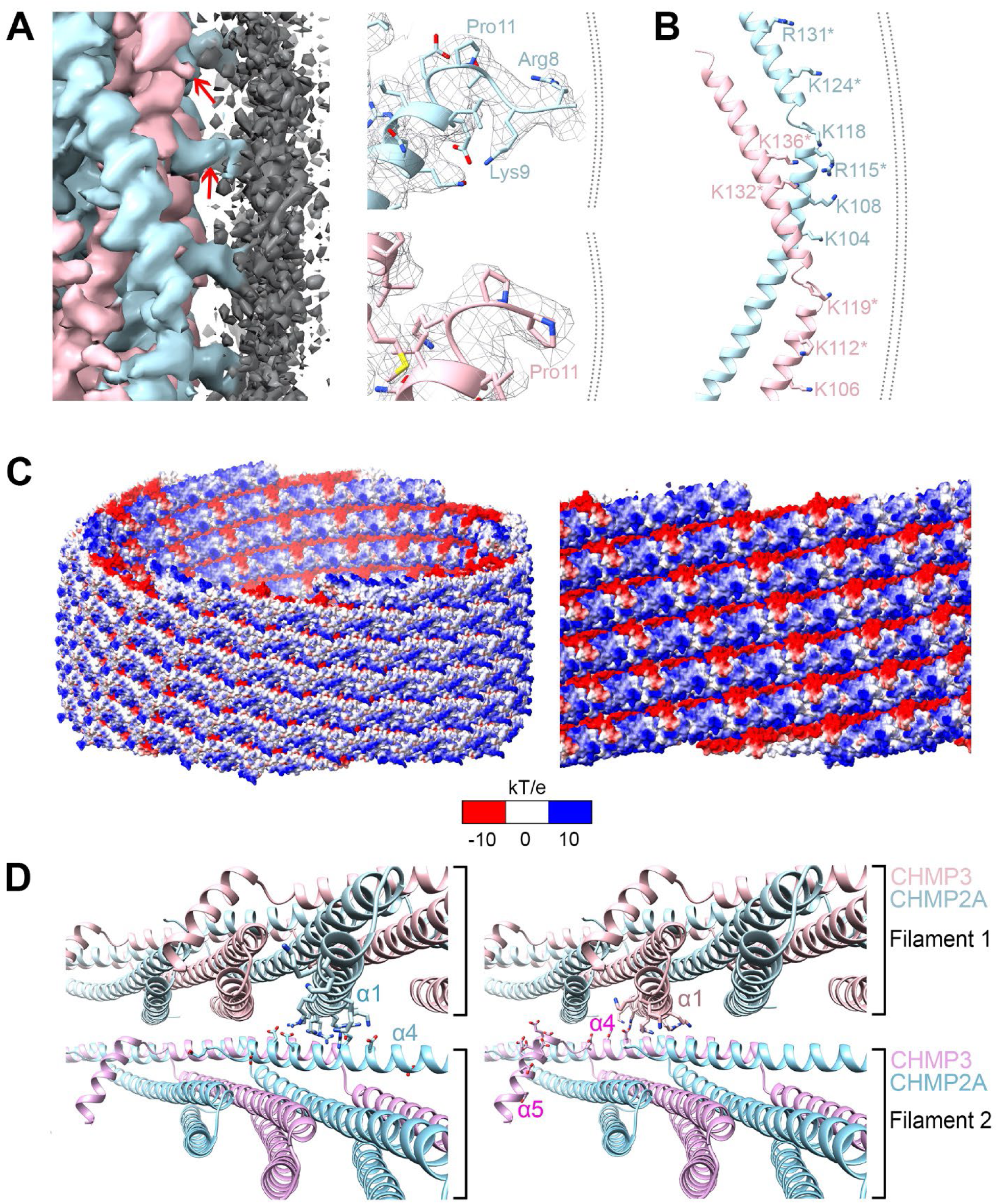
Membrane interaction of the CHMP2A-CHMP3 filament. **(A)** Left panel, zoomed-in view of the membrane-bound CHMP2A-CHMP3 filament, highlighting the interface between lipid membrane (dark grey) and CHMP2A (light blue) and CHMP3 (pink). Red arrows are pointing to the N-termini of both CHMP2A and CHMP3 that are oriented towards the lipid bilayer. Right panel, the orientation of the N-termini is determined by Pro11 of CHMP2A (light blue) and CHMP3 (pink) shown with the density map. **(B)** Ribbon diagram of the CHMP2A-CHMP3 heterodimer indicating the basic residues (sticks) oriented towards the membrane. Basic residues conserved in yeast Vps2 and Vps24 are marked by asterisks. **(C)** Electrostatic potential map of the CHMP2A-CHMP3 filament (left panel, tilted, side view), revealing the exposure of the cluster of basic charges, a small negatively charged surface and a neutral surface to the membrane. Right panel, zoomed-in view of the electrostatic surface of the inside of the polymer showing clusters of negative charges in one filament juxtaposed to positive charges of the neighboring filament. **(D)** Close-up of the inter-filament interactions. Ribbon diagram of two neighboring filaments showing the basic and acidic residues of CHMP2A (left panel) and CHMP3 (right panel) implicated in electrostatic inter-filament interactions.

### Inter-filament interactions

Conserved basic helix 1 residues of CHMP2A and CHMP3 (**Figure S6A**) are at the filament interface opposed by a stretch of conserved acidic residues within helices 4 and 5 (**Figure S6A**) of neighboring filaments (**Figure 3D**), which indicate electrostatic inter-filament interactions (**Figure 3C**). Mutation of the helix 1 basic cluster within either CHMP2A or CHMP3 prevented polymer formation *in vitro* (**Figure S7D**), indicating that the basic charge of helix 1 is important for filament polymerization, which is in line with mutagenesis of a similar cluster of basic residues within helix 1 of CHMP3 abolishing its dominant negative effect on HIV-1 budding (Muziol et al., 2006). To further test the electrostatic inter-filament interactions, we exposed the helical tubular CHMP2A-CHMP3 polymers to high ionic strength. This led to the partial unwinding of the filaments producing single and multi-stranded filaments (**Figure S8A-E**) in agreement with the presence of single and multi-start helices upon CHMP2A-CHMP3 polymerization *in vitro* (Effantin et al., 2013). We suggest that these electrostatic interactions between filaments enable filament sliding upon VPS4-catalyzed remodeling. The acidic cluster in helices 4 and 5 is conserved in CHMP4A, B, C, CHMP5 and CHMP6 (**Figure S6B**) indicating potential similar involvement in inter-filament interaction for the formation of mixed filaments. Notably, the acidic cluster is not conserved in CHMP1A and B, which stabilizes positively curved membranes via basic charges present on the inside of the protein tube-like polymer (McCullough et al., 2015). We therefore suggest that the acidic cluster is a hallmark of ESCRT-III stabilizing negatively curved membrane structures.

### VPS4 remodels and cleaves CHMP2A-CHMP3 membrane tubes

We next tested whether VPS4B can remodel the CHMP2A-CHMP3 membrane coated tubes as we have shown before for CHMP2A-CHMP3 tubes without membrane (Maity et al., 2019). When we incubated CHMP2A-CHMP3 and VPS4B containing membrane tubes (**Figure S9A**) with ATP and Mg^2+^, complete disassembly of the tubes was observed (**Figure S9B and C**). In order to image tube remodeling by fluorescence microscopy, VPS4B and caged ATP were incorporated into the tubes wrapped with fluorescently labelled membrane. Imaging of tubes containing either only caged ATP (**Figures 4A** **and** **B**; **movie 1; Figures S10A and B; movie 2)** or only VPS4B (**Figures 4C** **and** **D; movie 3**) demonstrated that photolysis used to uncage ATP did not affect the tube structure. However, tubes containing both caged ATP and VPS4B revealed constriction and tube cleavage upon ATP activation at different sites starting at 30s leading to complete disassembly within 270s (**Figures 4E** **and** **F; movie 4**) or starting at 69s (**Figures S10C and D; movie 5**) or 50 s (**movie 6**).

**Figure 4:**
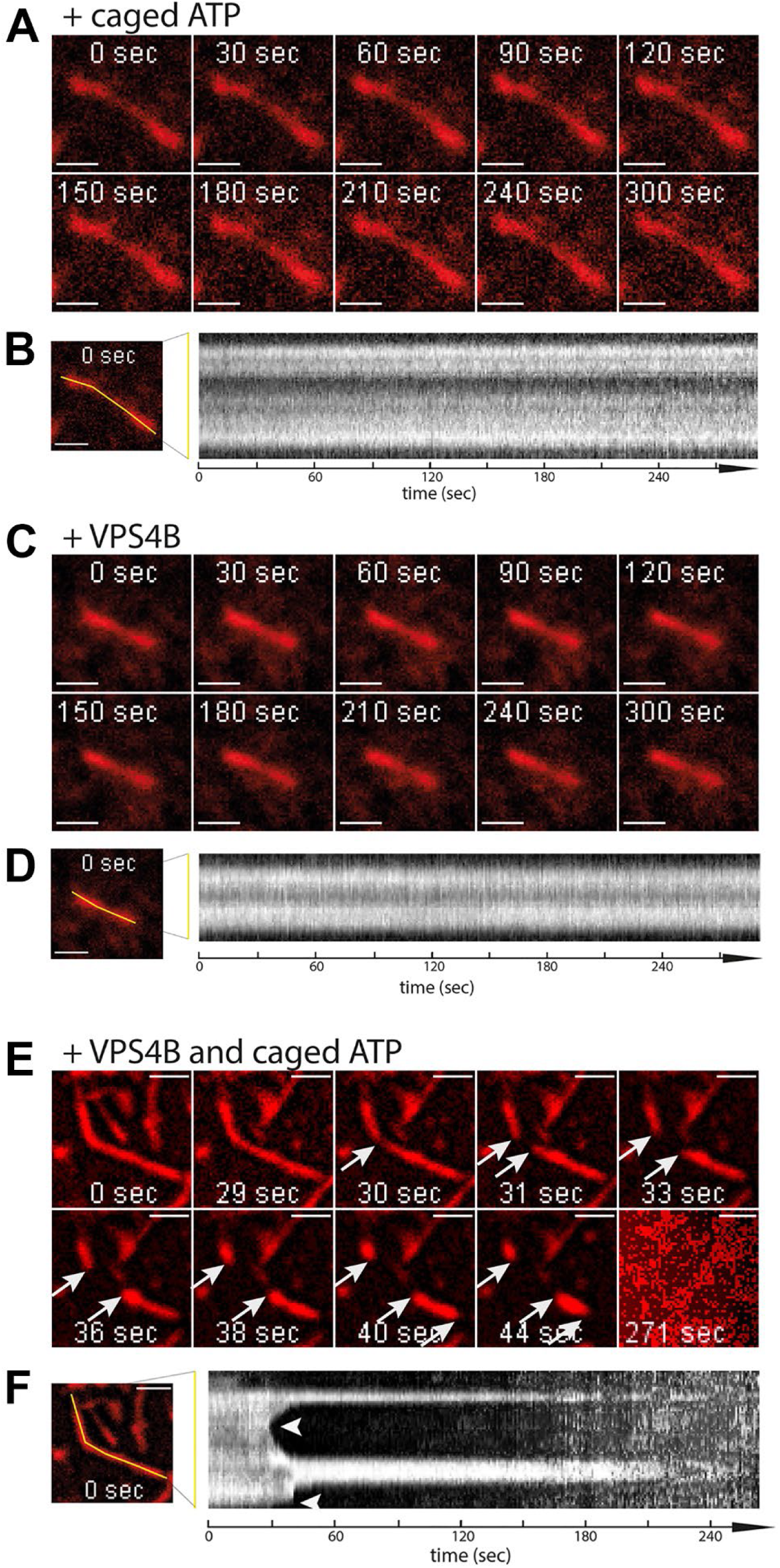
Imaging of VPS4B and ATP induced cleavage of CHMP2A-CHMP3 membrane coated tubes. **(A)** A CHMP2A-CHMP3-caged ATP containing membrane coated tube was activated at 365 nm (10%, 10s) to uncage ATP and imaged over 300s (movie S1). Scale bar, 1µm. **(B)** The kymograph of the tube shows that the tube stays intact over the imaging time. **(C)** A CHMP2A-CHMP3-VPS4B containing membrane coated tube was activated at 365 nm (10%, 10s) to uncage ATP and imaged over 300s. Scale bar, 1µm. **(D)** The kymograph of the tube shows that the tube stays intact over the imaging time. **(A)** to **(D)** demonstrate that imaging at 550 nm to visualize the membrane tube and ATP uncaging at 365 nm did not change the tube structures (movie S2). **(E)** Imaging of a CHMP2A-CHMP3-VPS4B-caged ATP containing membrane-coated tubes following ATP uncaging (365 mm, 10%, 10s) reveals constriction and cleavage of the tube at 30 s followed by a shrinking event from both sides. Another shrinking event is observed at 40s. Eventually all tubes were fully disassembled 271 s (movie S3 and S4). Scale bar, 1µm. **(F)** The kymograph of the tube indicates the kinetics of cleavage and shrinking.

Tube cleavage was further confirmed by high-speed AFM (HS-AFM) imaging. First, CHMP2A-CHMP3 tubes with and without membrane were imaged (**Figures S11A-C**). A comparative height histogram showed an increase in tube height of ∼8 nm for the membrane coated tubes (**Figure S11D**), as expected for an unilamellar membrane coating. Next, membrane-coated tubes loaded with 10 µM caged ATP with and without UV exposure (**Figure S11E; movie 7**) were recorded by HS-AFM. In the absence of VPS4B, photolysis of the caged ATP did not induce changes in tube morphology, consistent with the observations using fluorescence microscopy. Further, no changes in tube morphology were observed without UV exposure for an extended period of time of CHMP2A-CHMP3 tubes coated with membrane and loaded with VPS4B and caged ATP **(Figure 5A, movie 8)**. However, upon UV exposure, constriction and cleavage of the membrane coated tubes was observed (**Figure 5B; movie 9**). Kymographs along the tube cross section (**Figure 5C**) and the evolution of the height at the constriction sites over time (**Figure 5D**) reveal that complete cleavage of the membrane tube occurs within a time period of ∼ 200 s of UV exposure. We conclude that VPS4B can constrict the CHMP2A-CHMP3 filaments bound to membranes that leads to membrane cleavage, reminiscent of membrane fission.

**Figure 5.**
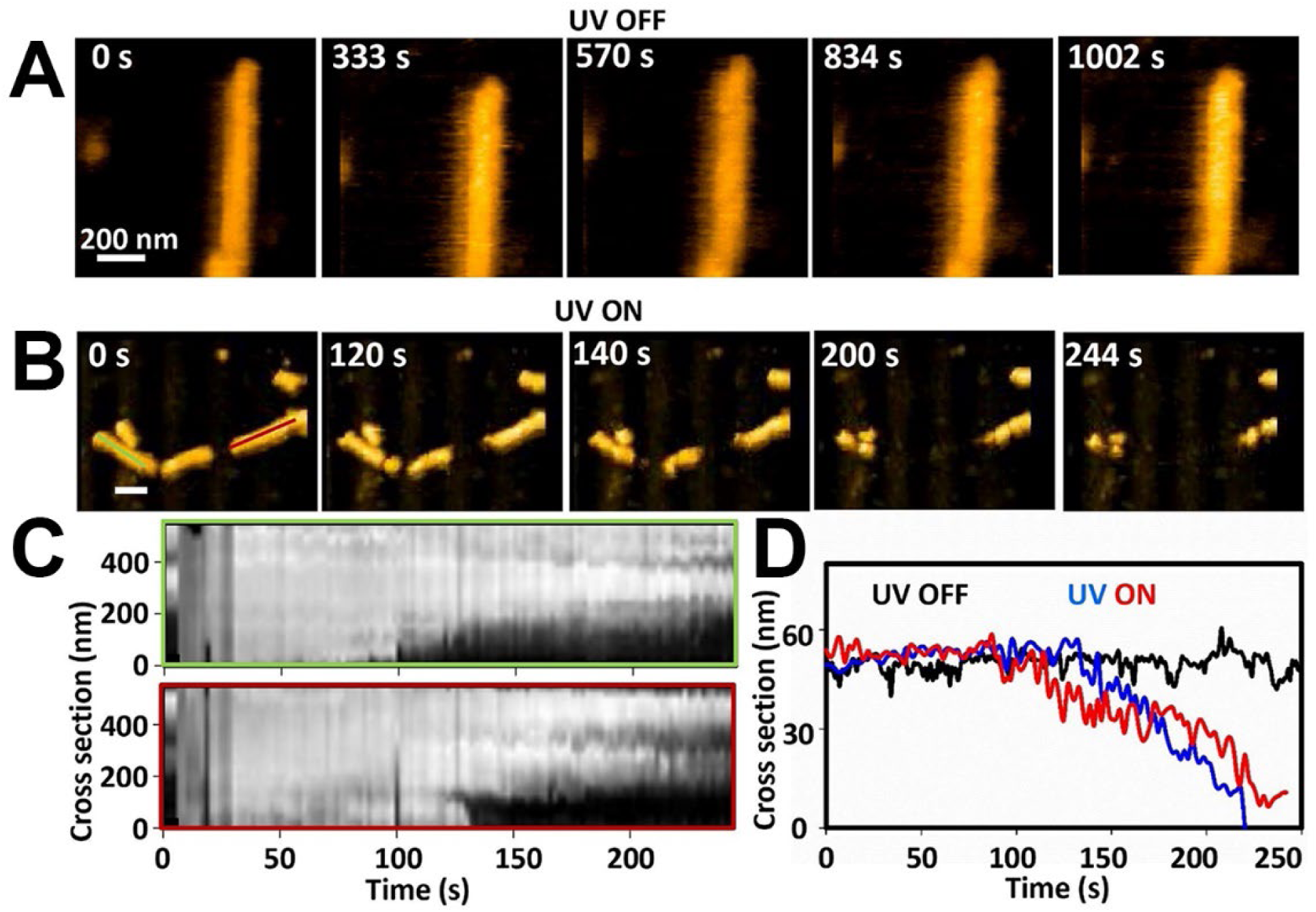
Constriction, cleavage, and disassembly of membrane coated CHMP2A-CHMP3 tube. **(A)** Snapshots of HS-AFM images of CHMP2A-CHMP3 tubes coated with membrane and loaded with 5 µM VPS4B, 10 mM caged ATP, in absence of UV irradiation. Scale bar, 200 nm. **(B)** As in A, but upon 365 nm UV irradiation. Scale bar, 200 nm. **(C)** Kymograph representation of the height vs time along the two lines in panel B (leftmost image) throughout the movie S10. **(D)** Example of height vs time profile of the membrane coated tube in absence of UV irradiation (in black), and in presence of UV irradiation (in blue and red).

## Discussion

The structure of the CHMP2A-CHMP3 heteropolymer demonstrates how ESCRT-III filaments polymerize into rigid structures that can stabilize and/or shape negatively curved membrane necks with diameters of approximately 50 nm. Such membrane structures are present at many ESCRT-catalyzed processes including vesicle and virus budding or at later stages during cytokinetic midbody constriction (McCullough et al., 2018; Vietri et al., 2020). Although the structural principles of the open ESCRT-III conformation are highly conserved between CHMP2A/CHMP3 and CHMP1B (McCullough et al., 2015), Snf7/Shrub (CHMP4) (McMillan et al., 2016; Tang et al., 2015) and bacterial PspA and Vipp1 (Gupta et al., 2021; Junglas et al., 2021; Liu et al., 2021), differences in helical hairpin stacking and orientations of the helical arms dictates the geometry of the filaments that stabilize positively curved membrane or negatively curved membrane as in case of CHMP2A-CHMP3 polymerization. Furthermore, assembly modes of ESCRT-III monomers can vary as shown for CHMP2A, CHMP3 and CHMP4B, which can yet adapt another filament geometry that stabilizes positively curved membranes (Bertin et al., 2020; Moser von Filseck et al., 2020). Thus the plastic nature of ESCRT-III protein conformations can lead to variable ESCRT-III filament geometries that can adapt a wide range of curvatures to accommodate ESCRT function in different membrane remodeling processes (Pfitzner et al., 2021).

A striking feature of the CHMP2A-CHMP3 polymer is the narrow range of tube diameters, indicative of a late recruitment during the constriction process. During yeast MVB biogenesis, Vps24/Vps2 (CHMP3/CHMP2) is indeed recruited last prior to complete ESCRT-III disassembly and probably completion of fission (Babst et al., 2002; Saksena et al., 2009; Teis et al., 2008). Likewise, CHMP4 isoforms recruit CHMP3 and CHMP2A to HIV-1 budding sites (Johnson et al., 2018; Morita et al., 2011; Prescher et al., 2015). The narrow range of diameters of the CHMP2A-CHMP3 polymer structure suggests that recruitment of CHMP3 and CHMP2A molds membrane necks into approximately 40 to 50 nm diameters, which sets the stage for further constriction. Comparison of the 410 and 430 Å wide structures shows that removal of only one CHMP2A-CHMP3 heterodimer per filament turn reduces the tube diameter by 20 Å, indicating how a stepwise removal of heterodimers can successively induce membrane constriction.

Another feature of the tubular polymer is that it can assemble from single or multi-stranded filaments (Effantin et al., 2013) the latter being the preferred assembly *in vitro*. Interaction between filaments is driven by complementary positive and negative charges. However, surprisingly, high ionic strength did not disassemble the tube consistently into individual filaments but partially preserved the multi-strand architecture *in vitro*. The 17.5 nm width of the six-stranded helix present in the structure fits the 17 nm wide helices imaged at the midbody (Guizetti et al., 2011), suggesting that such spirals contain six ESCRT-III filaments. Since inter-filament interactions are electrostatic, different ESCRT-III filaments may contribute to the formation of mixed multi-stranded filaments (Mierzwa et al., 2017; Pfitzner et al., 2020). In line, acidic residues within helix 4 and basic residues within helix 1 have been implicated in *S. cerevisiae* Snf7-Vps24 interaction (Banjade et al., 2019). Because basic and acidic charges within these regions are conserved in CHMP6, CHMP4A, B, C, CHMP2B and CHMP5, filaments thereof may also form side by side via homo- or hetero-filament assembly. The loose electrostatic inter-filament interactions likely facilitate sliding of filaments upon ESCRT-III filament remodeling by VPS4, which catalyzes filament constriction prior to complete disassembly (Maity et al., 2019). Notably, dynamic VPS4-mediated turnover of ESCRT-III has been proposed in different membrane remodeling processes (Adell et al., 2014; Adell et al., 2017; Mierzwa et al., 2017).

CHMP2A-CHMP3 polymers have been suggested to interact with negatively charged membranes (De Franceschi et al., 2018; Lin et al., 2005; Whitley et al., 2003), which is confirmed by the cluster of basic residues within the membrane interaction surface. Furthermore, the structure indicates that short N-terminal hydrophobic motifs, implicated in ESCRT-III function (Bodon et al., 2011) (Buchkovich et al., 2013) are positioned to insert into the membrane. Although this N-terminal motif is helical in a filament structure assembled by an intermediate Vps24 (CHMP3) conformation (Huber et al., 2020), the corresponding helices of CHMP2A and CHMP3 are not visible in the membrane-bound structure suggesting that the putative amphipathic helices can adopt different membrane insertion angles. Polymer interaction with membrane is tight, which excludes other membrane proteins thereby serving as a diffusion barrier (De Franceschi et al., 2018).

CHMP3 is dispensable for HIV-1 budding (Morita et al., 2011) and *S. cerevisiae* Vps24 can be substituted by Vps2 overexpression restoring partial endosomal cargo sorting (Banjade et al., 2021). Both processes depend on CHMP2A and Vps2 interaction with CHMP4 or Snf7 (Teis et al., 2008), (Morita et al., 2011). The structure suggests that CHMP3 can be structurally replaced by CHMP2A in the polymer, indicating that CHMP2A filaments on their own may form similar helices. Although CHMP2A can polymerize into circular filaments with approximate diameters of 40 nm that often coil up, no regular CHMP2A tube-like structures have been yet imaged *in vitro* (Effantin et al., 2013).

VPS4B remodels CHMP2A-CHMP3 helical tubular structures *in vitro* (Caillat et al., 2015; Lata et al., 2008b) inducing filament constriction and cleavage that generates dome-like end caps prior to complete disassembly, which was proposed to drive membrane fission (Fabrikant et al., 2009; Maity et al., 2019). Here, we show that VPS4B constricts and cleaves CHMP2A-CHMP3 membrane-coated tubes via membrane fission likely via the formation of dome-like end-caps (Maity et al., 2019). Cleavage of membrane tubes pulled from GUVs has been reported previously by employing a minimal system composed of *S. cerevisiae* Snf7, Vps24, Vps2 and Vps4 (Schoneberg et al., 2018), while another model proposed sequential recruitment of *S. cerevisiae* Snf7, followed by Vps2-Vps24, Vps2-Did2 and Did2-Ist1 for final constriction (Pfitzner et al., 2020). Our data suggest that CHMP2A-CHMP3 filaments constitute together with VPS4 a minimal ESCRT-III membrane fission machinery that can constrict membrane necks with 40 to 50 nm large diameters to the point of fission. It is yet unclear how many helical turns are required for constriction *in vivo*, although more than one, as estimated from imaging (Adell et al., 2017), would allow filament sliding powered by ATP-driven forces that can drive filament-induced membrane constriction and cleavage. Finally, catalyzing membrane fission with a minimal machinery is well in line with ancestral ESCRT-III function (Caspi and Dekker, 2018; Ithurbide et al., 2022).

## Methods

### Expression and purification

CHMP2AΔC containing residues 1 to 161 was subcloned in a pMAL-c5X vector with an additional TEV site at the amino terminal end and expressed as a MBP fusion protein in the C41 (DE3) *E. coli* bacterial strain (Lucigen). Expression was induced for 1h at 37°C. Bacteria were lysed by sonication in a buffer containing 50 mM HEPES pH 7.5, 300 mM NaCl, 1 mM DTT, 5 mM EDTA and protease inhibitors. Cleared lysate was applied onto an amylose resin (New England Biolabs), washed with buffer A (25mM HEPES 7.5, 150 mM NaCl, 1mM DTT), then with Buffer B (25 mM HEPES pH 7.5, 1 M NaCl, 1 M KCl, 1mM DTT) followed by a last wash with Buffer A. Finally, protein was eluted with buffer C (25 mM HEPES 7.5, 150 mM NaCl, 10 mM maltose). The most concentrated fraction was directly applied to size exclusion chromatography (SEC) Superdex 200 column (GE Healthcare) in buffer D (25 mM HEPES pH 7.5, 150 mM NaCl).

Full-length CHMP3 was subcloned in a pProEX-HTb vector (Life Technologies, Thermo Fisher) and expressed in BL21Gold (DE3) *E. coli* bacterial strain (Agilent). Expression was induced for 3h at 37°C and purified as described (Lata et al., 2008b) with minor modifications. Bacteria were lysed by sonication in buffer E (25 mM HEPES pH 7.5, 150 mM NaCl, 10mM imidazole) containing protease inhibitors and the cleared lysate was applied onto a Ni^2+^-chelating sepharose (Cytivia), washed extensively with lysis buffer E, and subsequently with buffer F (25 mM HEPES pH 7.5, 300 mM NaCl, 300 mM KCl, 20 mM imidazole) and buffer G (25 mM HEPES pH 7.5, 300 mM NaCl, 300 mM KCl, 50 mM imidazole). Finally, CHMP3 was eluted with buffer H (25 mM HEPES 7.5, 150 mM NaCl, 350 mM imidazole) and cleaved overnight at 4°C with Tobacco Etch Virus (TEV) protease at 1:100 (w/w) ratio in the presence of 10 mM β-mercaptoethanol. Cleaved protein was then applied on a second Ni^2+^-chelating sepharose in order to remove TEV, the His-tag and uncleaved protein. The final step included size exclusion chromatography (SEC) on a Superdex 75 column (GE Healthcare) in buffer D. CHMP3 concentrated at 300 µM was frozen for further use.

CHMP2AΔC-mutants containing residues 9 to 161 and CHMP3-mutants containing residues 9 to 183 were synthesized (ThermoFisher), subcloned in a pETM40 vector (PEPcore facility-EMBL Heidelberg) and the pProEX-HTb (ThermoFisher) vector, respectively. Mutants were expressed and purified as described above for wild type sequences.

### CHMP2A-CHMP3 membrane tube generation

For helical tube formation as described previously (Lata et al., 2008b), 10 µM CHMP2AΔC was mixed with 20 µM full-length CHMP3 and incubated for 48-72h at 4°C. After incubation, tubes were harvested by centrifugation at 20,000g for 30min and the pellet containing CHMP2AΔC-CHMP3 tubes was resuspended in buffer D. In order to wrap the CHMP2AΔC-CHMP3 tubes with a lipid bilayer, the following lipid film was produced containing 70% Egg phosphatidyl choline (Egg PC), 10% dioleoyl glycero phosphoserine (DOPS), 10% dioleoyl glycerol phosphoethanolamine (DOPE), 10% brain phosphatidylinositol-4,5-bisphosphate (PI(4,5)P2) and 2µL of dioleoyl-sn-glycero-3-phosphoethanolamine-N-(lissamine rhodamine B sulfonyl) (LISS-Rhodamin PE) (all Avanti Polar lipids). The lipid film was resuspended in water at a final concentration of 5 mg/mL. The CHMP2AΔC-CHMP3 tubes (25 µL) were mixed with 25 µL of 2% CHAPS, 25 µL of lipids and 0.1 mg/mL of TEV protease (to remove the MBP from CHMP2AΔC) and incubated at room temperature for 2h. To remove free lipids/micelles and CHAPS, the tubes were dialyzed twice for 48-72h against buffer I (25mM Tris pH 7.4, 25 mM NaCl, 1 mM β-mercaptoethanol and 0.5 g of Bio-Beads (Biorad). After dialysis, CHMP2AΔC-CHMP3 tubes wrapped with bilayer were incubated with Bio-Beads overnight at 4°C and removed by centrifugation. The quality of the bilayer wrapped CHMP2AΔC-CHMP3 tubes was assessed by negative staining EM prior to cryo-EM data collection, fluorescence microscopy imaging and HS-AFM analysis.

To test remodeling by VPS4, CHMP2A-CHMP3 tubes were incubated with 10 to 20 µM VPS4B, 5 mM MgCl_2_ and 5 or 10 mM caged ATP (#A1048 Invitrogen) prior to membrane wrapping, following the protocol described above. Because deposition of the membrane onto the CHMP2A-CHMP3 protein coat requires extensive dialysis, the final VPS4 concentration present within the tubes can be only estimated from SDS-PAGE; however, the final concentration of caged ATP inserted into the tubes cannot be determined.

### CHMP2A-CHMP3-VPS4 membrane tube imaging

Epifluorescence video microscopy of CHMP2A-CHMP3 membrane tubes containing VPS4B and caged ATP was performed using an Olympus IX83 optical microscope equipped with a UPFLN 100X O-2PH/1.3 objective and an ORCA-Flash4.0 Digital sCMOS camera (Hamamatsu). A 5 μL aliquot of ESCRT-III tube suspension was spread on a slide, covered with a glass coverslip (#1) and sealed with twinsil speed 22 (Picodent, ref 13001002) for imaging. Caged-ATP was uncaged using a 10s 10% 365-nm LED illumination (**Figures 4A-F** **and movies S1-4**) or using at each time point a 100ms 30% 365-nm LED illumination (**Figure S10 and movies S5 and S6**). ESCRT tubes were fluorescently imaged using a 550nm LED (10% with an exposure time of 100ms) at 1 frame/s. Images were acquired using the Volocity software package. Images were analyzed, adjusted, and cropped using ImageJ software.

### HS-AFM analysis

The AFM images were acquired in amplitude modulation tapping mode in liquid, using high-speed atomic force microscopes (RIBM, Japan) (Maity et al., 2019; Maity et al., 2020). The HS-AFM imaging was performed using USC-F1.2-k0.15 cantilevers (NanoWorld, Switzerland), an ultrashort cantilever with a nominal spring constant of 0.15 N/m and a resonance frequency ≈ 0.5 MHz. All HS-AFM recordings were done at room temperature and in buffer D. Uncaging of the caged ATP was performed by directly irradiating 365 nm UV light at the AFM sample stage using an optical fiber. The membrane coated tubes were immobilized at the surface using streptavidin on top of a lipid bilayer (DOPC) on mica containing 0.01% biotinylated lipid (Keya et al., 2017). HS-AFM images were analyzed using Igor Pro, and ImageJ with additional home written plugins (Maity et al., 2022). Height measurements were performed on raw images after tilt correction.

### Dominant negative effect of CHMP3(1-150) wild type and mutants

CHMP3 residues 1-150 wt or mutants were synthesized (ThermoFisher) and cloned into the pEGFP-N1 vector using restriction sites *Xho*1-*Hind*III. To determine the effect of GFP-CHMP3 (1-150) on virus like particle (VLP) production upon HIV-1 Gag expression, 293T cells were seeded into 10mm dishes and transfected 24 hr later using a Jetprime (Polyplus) technique. The cultures were co-transfected with 0.5 μg of Rev-independent HIV-1 Gag construct and with 2 μg of either pcDNA or wild type and mutant versions of GFP-CHMP3(1-150). Twenty-four hours post transfection, VLPs released into the culture medium were pelleted through sucrose. HIV-1 Gag proteins in VLPs and cell lysates were detected by Western blotting with a mouse anti-p24 antibody (183-H12-5C). For life cell imaging cells were seeded in glass bottomed μ-dishes and co-transfected with 0.8 μg of Rev-independent HIV-1 Gag construct, 0.2 μg of Gag-mCherry (Jouvenet et al., 2008) and with 1 μg of either wild type or mutant GFP-CHMP3(1-150). ESCRT-III and Gag protein localization was analyzed by spinning disc confocal microscopy 24 hr post transfection in HeLA cells.

### Cryo-EM sample preparation and data collection

*Cryo-electron microscopy.* 3.5 µL of sample were applied to glow discharged (45s 30 mA) 1.2/1.3 Ultrafoil holey grids (Quantifoil Micro Tools GmbH, Germany) and they were plunged frozen in liquid ethane with a Vitrobot Mark IV (Thermo Fisher Scientific) (100% humidity, temperature 20°C, 6 s blot time, blot force 0). The grids were pre-screened on the 200kV Glacios electron microscope (Thermo Fischer Scientific) at the IBS (Grenoble) and data were collected at the beamline CM01 of the ESRF (Grenoble, France) (Kandiah et al., 2019) on a Titan Krios G3 (Thermo Fischer Scientific) at 300 kV equipped with an energy filter (Bioquantum LS/967, Gatan Inc, USA) (slit width of 20 eV). 5028 movies were recorded automatically on a K2 summit direct detector (Gatan Inc., USA) with EPU (Thermo Fisher Scientific) for a total exposure time of 5 s and 200 ms per frame resulting in 25 frame movies with a total dose of ∼24 e^−^/Å^2^. The magnification was 130,000x (1.052 Å/pixel at the camera level). The defocus of the images was adjusted between −0.5 and −1.5 μm in 0.2 μm steps. For the high ionic strength unwinding of the CHMP2A-CHMPA filament the same grid and freezing conditions have been used as described above and images have been recorded on the Glacios electron microscope using a K2 direct electron detector.

### EM image analysis and 3D reconstructions

The workflow of the image analysis is shown in **Figure S2**. Electron beam-induced sample motion on the recorded movie frames was corrected using MotionCor2 (Zheng et al., 2017) and the contrast transfer function (CTF) was estimated with CTFFIND4 (Rohou and Grigorieff, 2015). 9,207 filaments were manually picked from 5,027 micrographs using the EMAN2 program e2helixboxer.py (Tang et al., 2007). All subsequent data processing steps were carried out in RELION3 (Scheres, 2012; Zivanov et al., 2018) unless mentioned otherwise. Initially, 89,122 overlapping segments were extracted with ∼90% overlap between boxes of 768 x 768 pixels and down sampled to a pixel size of 2.104 for initial classification steps. Several rounds of 2D classification resulted in class averages that could be classified into 5 main different groups based on the filament diameter, without membrane (380 Å diameter (7.4%), 410 Å diameter (51.4%), 430 Å diameter (31.2%), 470 Å diameter (1.9%) and 490 Å diameter (0.3%)). In order to compensate for potential mis-assignment of diameters to the segments due to inaccuracies in 2D classification, we assigned to each entire tube a diameter based on the class assignment of the corresponding segments. If more than 80% of segments of a particular tube were belonging to classes assigned to a particular diameter, this entire tube would be assigned this diameter for the subsequent steps. After re-extraction of segments with ∼95% overlap, another round of 2D classification was performed for each diameter group. Most populated 2D classes with filament diameter of 430 Å and 410 Å were chosen for further processing and analysis. For determination of helical symmetry, the sum of power spectra from a smaller subset (1,904 and 3,993 segments from one 2D class each for 430 Å and 410 Å respectively) was calculated for both filament groups. The resulting average power spectrum (**Figures S1C and D**) was analyzed for estimation of helical symmetry parameters using the web tool helixplorer (http://rico.ibs.fr/helixplorer/). Based on a prior visual inspection of the PS, we made following hypotheses: the layer line with a maximum seemingly on the meridian could be the helical rise or the pitch (given the large diameter of the tube and possibilities that selected 2D class averages contained a number of slightly out-of-plane tilted segments). Given those two hypotheses, and allowing any cyclic symmetries, we explored possible helical symmetries matching the experimental PS, giving a list of 20 and 15 symmetries to test for the 430 Å and 410 Å diameter classes, respectively. Those symmetries were applied on the real-space 2D class-averages using SPRING program segclassreconstruct.py (Desfosses et al., 2014) in order to generate initial models and narrowing down possible symmetry solutions to 14 and 10, by discarding those giving aberrant density distribution. Using those initial models, each of the remaining symmetry solutions was tested for 3D refinement in RELION3, and the resulting maps inspected for high-resolution features such as clear secondary structures, allowing to determine the helical parameters to be 18 Å pitch, 2.72 Å rise, 54.39° twist, 6.6 units/turn and C2 point symmetry for 430 Å, and 9 Å pitch, 1.43 Å rise, 57° twist, 6.3 units/turn and C1 point symmetry for 410 Å diameter filaments.

In order to select a more homogeneous subset of segments, we applied 3D classification and the classes (containing 25,353 and 11,396 segments for diameters 430 and 410 Å) were chosen for a final round of 3D auto-refine reconstruction that converged to a 2.74 Å rise and 54.37° twist for 430 Å, and 1.44 Å rise and 57.04° twist for 410 Å diameter filaments. Using soft protein-only masks, the final resolutions were estimated at 3.3 Å and 3.6 Å for the 430 Å and 410 Å diameter filaments, at the FSC (Fourier shell correlation) 0.143 cutoff criterion (Rosenthal and Henderson, 2003). The maps were sharpened with bfactors of -96.57 Å^2^ (430 Å) and -101.52 Å^2^ (410 Å). Local resolution was estimated in RELION3 (Scheres, 2012) and the density maps were rendered in UCSF Chimera (Pettersen et al., 2004). The statistics of the EM map are summarized in **Table S1**.

### Atomic modelling and validation

The SWISS-MODEL (Waterhouse et al., 2018) server was used to create homology models of human CHMP3 and CHMP2A, using the open conformation of CHMP1B (PDB ID 6TZ4) as a reference model. Helices with residues 15-52, 57-117 and 120-151 for CHMP3 and 14-51, 56-116 and 119-150 for CHMP2A were initially fit into the EM density as separate rigid bodies using Chimera and then adjusted in Coot (Emsley et al., 2010). Further, the rest of the N-terminal and C-terminal residues, as well as the connecting loops were manually built and adjusted in Coot. The CHMP2A-CHMP3 heterodimer model was then expanded by helical symmetry in each direction in order to get 10 such dimers surrounding the central dimer. Thus, a total of 11 helical symmetry-related dimers were again checked in Coot, before applying the first round of real-space refinement in PHENIX (Adams et al., 2010) with non-crystallographic symmetry (NCS) restraints. NCS, along with SS (secondary structure) restraints were then used for a second round of real-space refinement. At last, the symmetry-related dimers were removed and the central CHMP2A-CHMP3 dimer was saved as the final model. The statistics of the final models were tabulated using MolProbity (Williams et al., 2018) and are summarized in **Table S1** and map versus atomic model FSC plots were computed in PHENIX (Afonine et al., 2018). All structure figures were generated with UCSF Chimera, ChimeraX (Goddard et al., 2018) and PyMOL (W. Delano; The PyMOL Molecular Graphics System, Version 1.8 Schrödinger, LLC, http://www.pymol.org). Sequence alignments were performed with Clustal Omega (Goujon et al., 2010) and ESPript (Robert and Gouet, 2014).

## Data Availability

Cryo-EM maps and models were deposited to the PDB and EMDB with the following codes: membrane-bound CHMP2A-CHMP3, 430 Å diameter (PDB ID 7ZCG, EMD-14630) and membrane-bound CHMP2A-CHMP3, 410 Å diameter (PDB ID 7ZCH, EMD-14631).

## Acknowledgement

This research was funded by the ANR (ANR-14-CE09-0003-01; ANR-19-CE11-0002-02)(W.W.). WW acknowledges support from the Institut Universitaire de France (IUF) and access to the platforms of the Grenoble Instruct-ERIC center (IBS and ISBG; UAR 3518 CNRS-CEA-UGA-EMBL) within the Grenoble Partnership for Structural Biology (PSB), with support from FRISBI (ANR-10-INBS-05-02) and GRAL, a project of the University Grenoble Alpes graduate school (Ecoles Universitaires de Recherche) CBH-EUR-GS (ANR-17-EURE-0003). The IBS electron microscope facility is supported by the Auvergne-Rhône-Alpes Region, the Fondation pour la Recherche Medicale (FRM), the FEDER/ERDF fund (European Regional Development Fund) and the GIS-IBiSA (Infrastructures en Biologie, Sante et Agronomie). We acknowledge the provision of in-house experimental time from the CM01 facility at the ESRF and we thank Leandro Estrozi for extensive discussion and help with helical image analysis.

## Author contributions

W.W. conceived the study, designed experiments, interpreted experiments, supervised and received funding for the study. K.A. performed all cryo-EM data analyses. N.D.F. established the membrane coating protocol. D.G. prepared wild type and mutant CHMP2A-CHMP3 polymers for all analyses. G.Su. (G. Sulbaran) performed negative staining EM analyses. C.B. and J.P.K. performed fluorescence microscopy imaging and H.W. mutant analyses. S.M. performed AFM analyses and W.H.R. supervised AFM analyses. G.E. and G.S. collected cryoEM data and A.D. supervised all aspects of cryoEM data analyses, structure solution and interpretation. W.W. wrote the paper with input from all authors.

## Supplemental Figures

**Figure S1:**
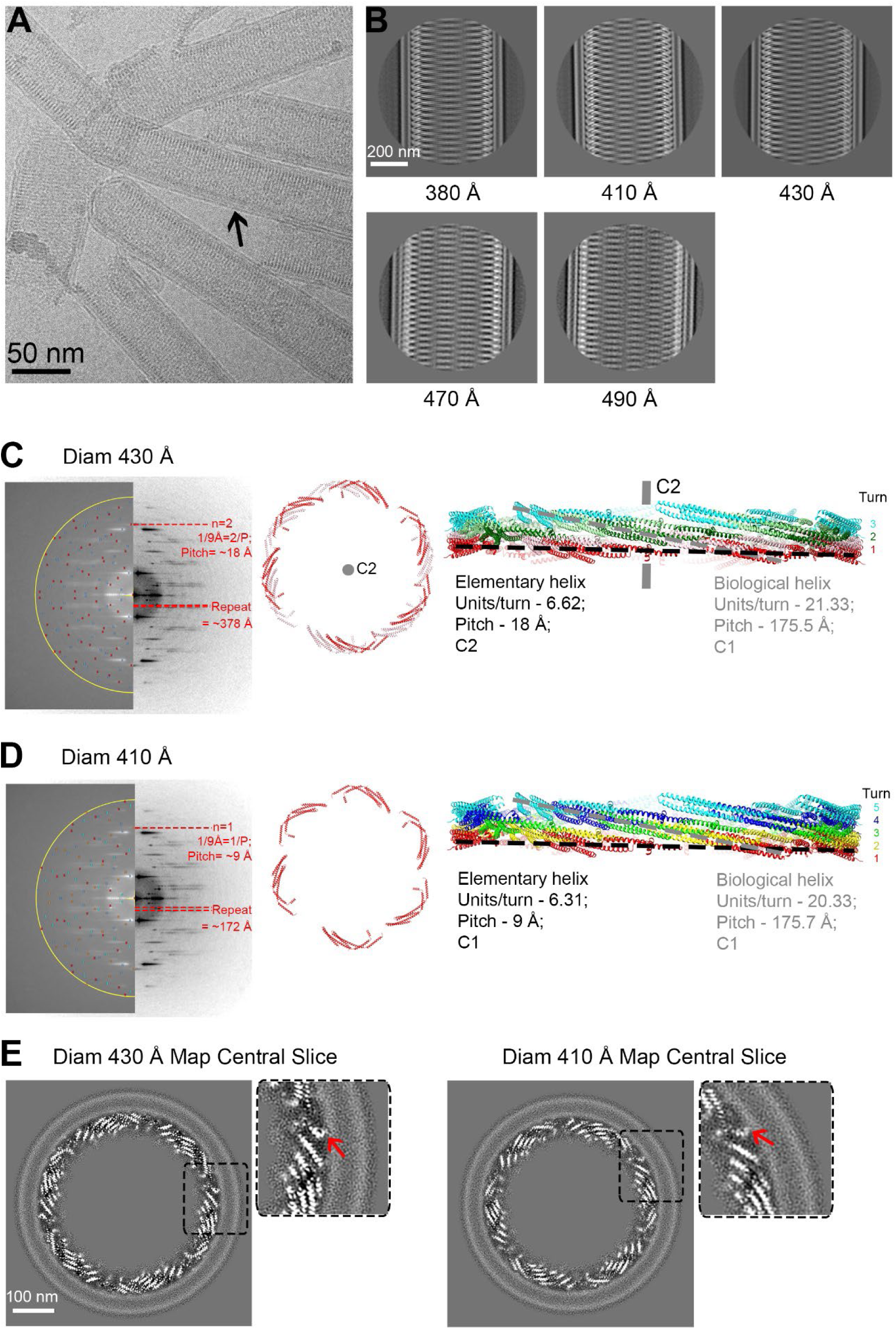
Cryo-EM data processing of CHMP2A-CHMP3 membrane-coated tubes and helical symmetry analyses. **(A)** Representative cryo-electron micrograph of CHMP2A-CHMP3 membrane tubes, with an arrow pointing to the lipid bilayer. Scale bar, 50 nm. **(B)** Selected 2D class averages of manually picked datasets, arranged according to the tube diameter ranging from 380 to 490 Å as indicated. **(C), (D)** Helical symmetry determination and representation of the elementary and biological helices for the 430 Å and 410 Å diameter tubes. Left panel, the sum of the 2D power spectra of segments corresponding to one class-average show in both cases a maximum on or near the meridian corresponding to the pitch (Bessel order n=1 for C1 helix; n=2 for C2 helix) instead of the axial rise, due to the large helical diameter and possible inclusion of slightly out-of-plane tilted segments in the class-average. The left half of the sum of the power spectra show the calculated position from helixplorer of the two first maxima of each Bessel function corresponding to the determined symmetry. Middle panel, six asymmetric units (CHMP2A-CHMP3 dimers; in red) of the elementary helix are represented. The symmetry related protomers of the 430 Å diameter are colored in light red. **(C**), (**D)** Right panel, side view of the elementary helix with turns (red, yellow, green, blue and aqua) indicated with a black dashed line. The grey dashed line follows one turn of the biological helix. The central grey line (in C) highlights the C2 symmetry axis. Symmetry parameters of both the elementary and biological helices are indicated. **(E)** Central slices looking down the helical axis of the cryo-EM 3D reconstructions of the 430 and 410 Å diameter tubes. The red arrows in the right zoom-in images indicate the density of the N-terminal region prone to insert into the lipid bilayer.

**Figure S2:**
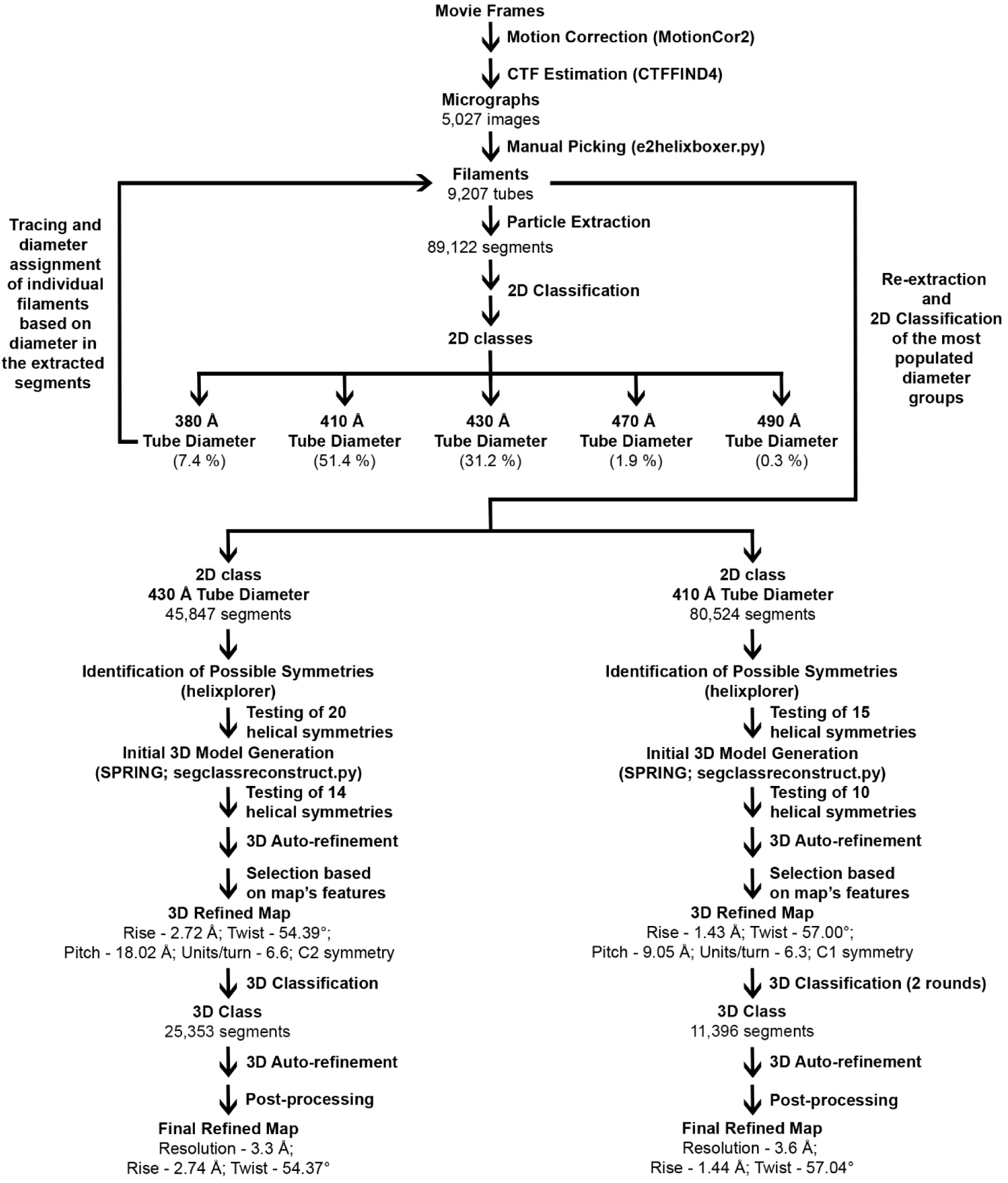
Cryo-EM image processing workflow of 430 and 410 Å diameter tubes structure determination. Basic image processing strategy used for helical 3D reconstruction and refinement of 430 and 410 Å diameter tubes is shown. Helical filaments were segmented and classified based on the tube diameter. Segment subsets were subjected to symmetry determination (http://rico.ibs.fr/helixplorer/) and initial 3D model generation in SPRING, followed by symmetry refinement and final 3D structure refinement in RELION. A complete description of the processing workflow is provided in ‘Materials and Methods’ section.

**Figure S3:**
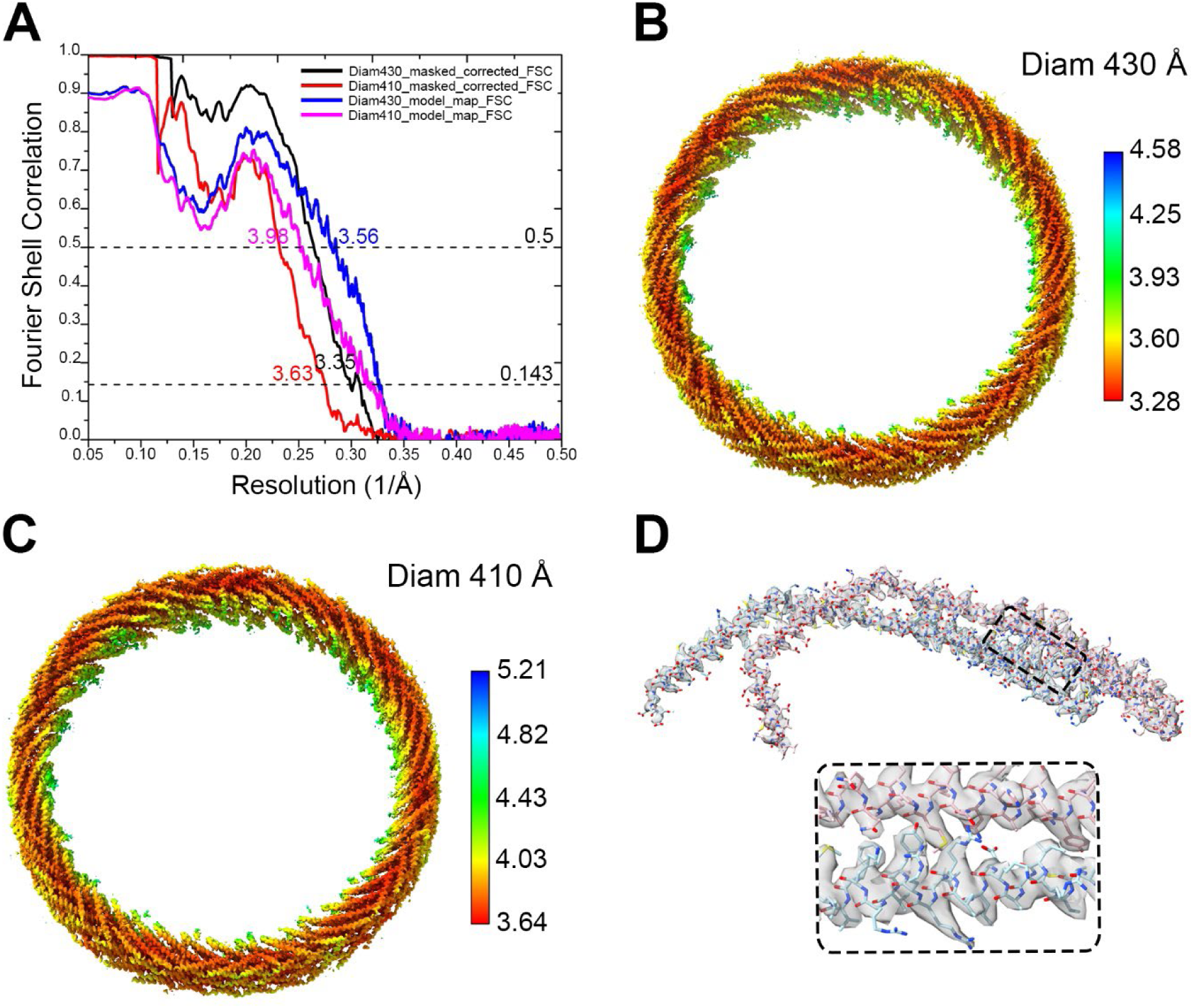
430 and 410 Å diameter tubes FSC curves, local resolution maps and atomic model fitting. **(A)** FSC curves for the 430 Å (black) and 410 Å (red) diameter tube maps, with the resolutions at the FSC cut-off of 0.143 are indicated. Model versus map FSC curves, with the resolutions at the FSC cut-off of 0.5 are indicated for the 430 Å (blue) and 410 Å (pink) diameter tube maps. Local resolution estimates are mapped onto the 430 Å **(B)**, and 410 Å **(C)** diameter tube cryo-EM density maps and the color keys (right) highlight the local resolution values in Å. **(D)** The refined atomic model of CHMP2A-CHMP3 dimer was fit into the corresponding cryo-EM density map of the 430 Å diameter tube. The inset (below) represents the zoomed-in view of the fitted model, indicating CHMP2A and CHMP3 helices and the corresponding map.

**Figure S4:**
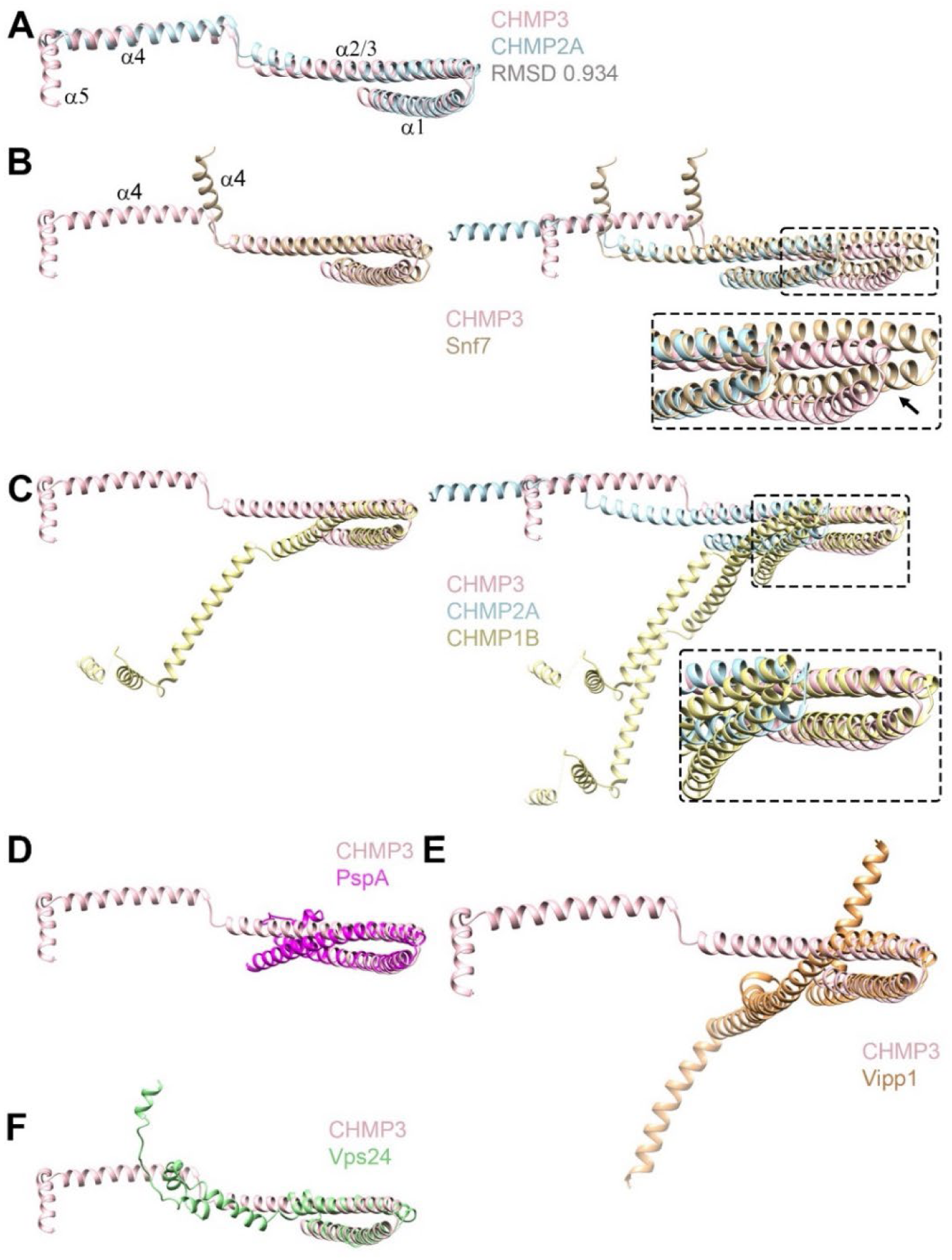
Comparison of ESCRT-III open conformations highlights their versatile polymerization modes. **(A)** Ribbon diagram of Cα superposition of CHMP2A and CHMP3 reveals an RMSD of 0.934 Å. **(B)** Ribbon diagram of Cα superposition of CHMP3 with an Snf7 monomer (left panel) and with the homodimer (crystallographic dimer) (right panel), which indicates different hairpin interaction (arrow) for the second protomer and different orientations of Snf7 helix 4. **(C)** Ribbon diagram of Cα superposition of CHMP3 and CHMP1B (left panel) and the superposition of the CHMP2A-CHMP3 heterodimer onto the CHMP1B homodimer (right panel) indicate the differences in helical hairpin stacking (zoom, right panel) and orientations of the C-terminal helical arms (helices 3 to 5). **(D)** Ribbon diagram of Cα superposition of CHMP3 (pink) with PspA and **(E)** with Vipp1. **(F)** Ribbon diagram of Cα superposition of CHMP3 with an intermediate Vps24 conformation that forms filaments on its own.

**Figure S5:**
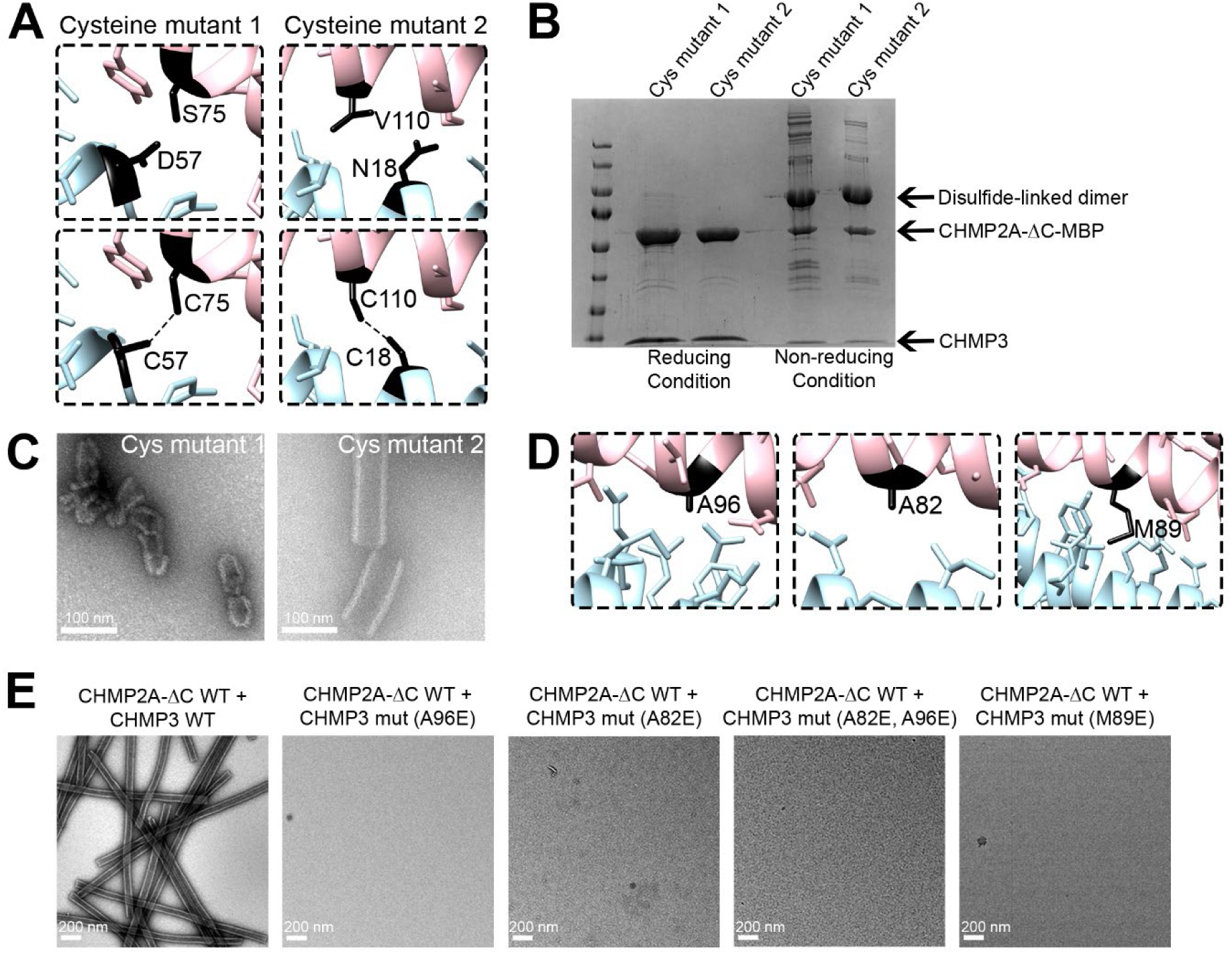
Structure-based mutagenesis of CHMP2A-CHMP3 heterodimer formation and polymerization *in vitro*. **(A)** Close-up views of the pairs of residues (black) mutated to cysteine to induce the formation of disulfide-linked CHMP2A (light blue) - CHMP3 (pink) heterodimers upon polymerization. **(B)** Cysteine cross-linking of the CHMP2A-CHMP3 heterodimer. Mutant CHMP2A_D57C was incubated with CHMP3_S75C and CHMP2A_N18C with CHMP3_V110C to induce polymerization as reported for wild-type CHMP2A and CHMP3 (Lata et al., 2008b). SDS-PAGE analysis showing that both CHMP2A_D57C-CHMP3_S75C and CHMP2A_N18C-CHMP3_V110C formed disulfide-linked dimers under non-reducing SDS PAGE conditions. **(C)** Negative staining electron micrographs showing regular tube formation for CHMP2A_N18C-CHMP3_V110C (right), while CHMP2A_D57C-CHMP3_S75C (left) produced only shorter tubes. Scale bar, 100 nm. **(D)** Close-up views of the CHMP3 interface residues A96, A82 and M89E tested for heterodimer formation and polymerization. **(E)** Negative staining electron micrographs of CHMP2A-CHMP3 wild-type and mutants (CHMP3_A96E, CHMP3_A82E, CHMP3-A82E_A96E and CHMP3_M89E) assemblies as indicated. Scale bar, 200 nm.

**Figure S6:**
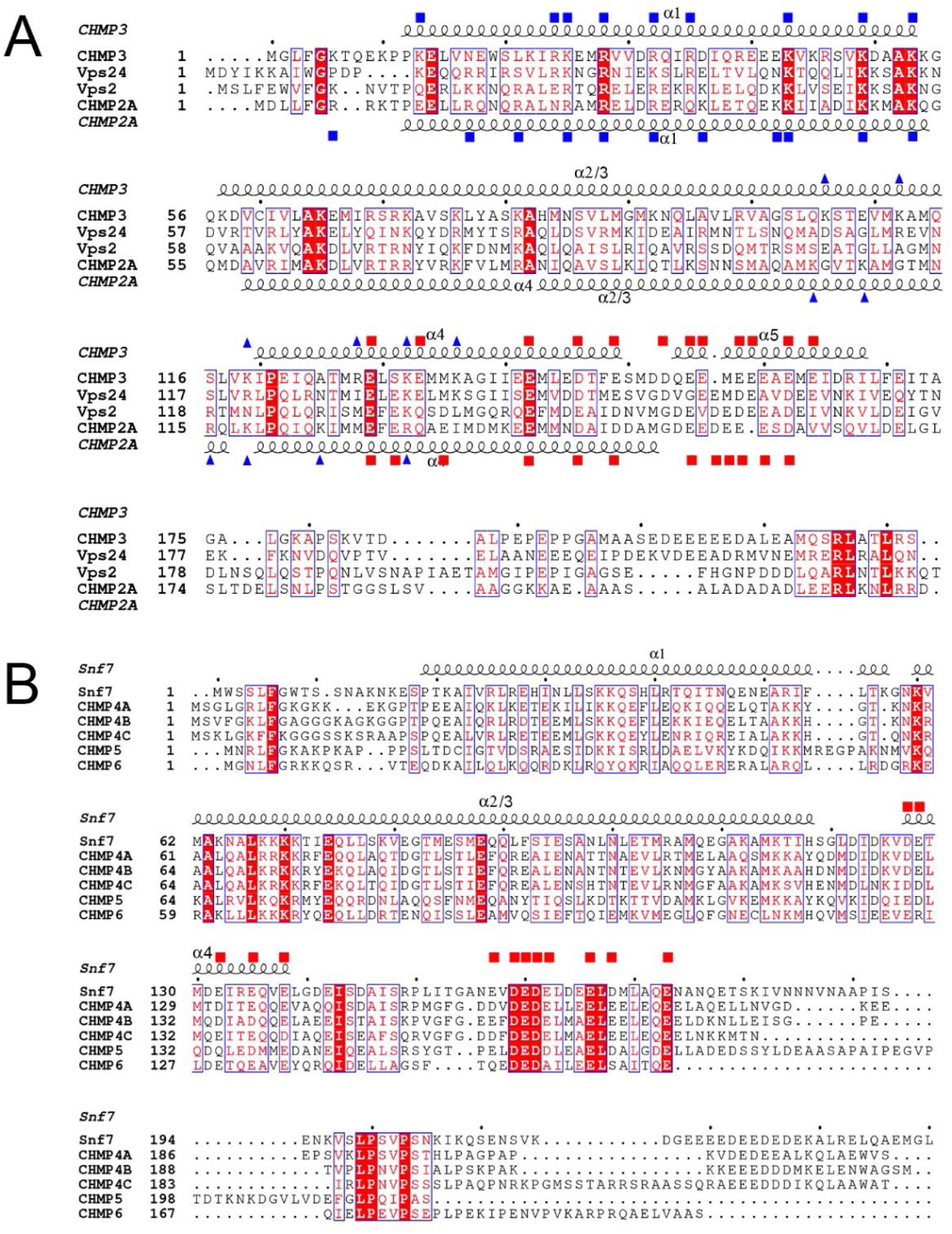
ESCRT-III sequence alignment. **(A)** Sequence alignment of CHMP3 (AF219226), *S. cerevisiae* Vps24 (QHB09957), *S. cerevisiae* Vps2 (P36108.2) and CHMP2A (NM_198426.3). Secondary structure elements are shown for CHMP3 above the sequence and for CHMP2A below the sequence alignment. Blue triangles indicate basic residues of CHMP3 (above) and CHMP2A (below) exposed at the membrane binding interface. Blue rectangles show basic residues and red squares conserved acidic residues exposed at the interface between filaments. **(B)** Sequence alignment of *S. cerevisiae* Snf7 (Z73197.1) and its secondary structure (pdb 5FD9), CHMP4A (NM_014169.5), CHMP4B (NM_176812.5) CHMP4C (NM_152284), CHMP5 (NM_016410.6) and CHMP6 (NM_024591.5). Conserved acidic residues implicated in inter-filament interactions in the CHMP2A-CHMP3 polymer are indicated as red squares.

**Figure S7:**
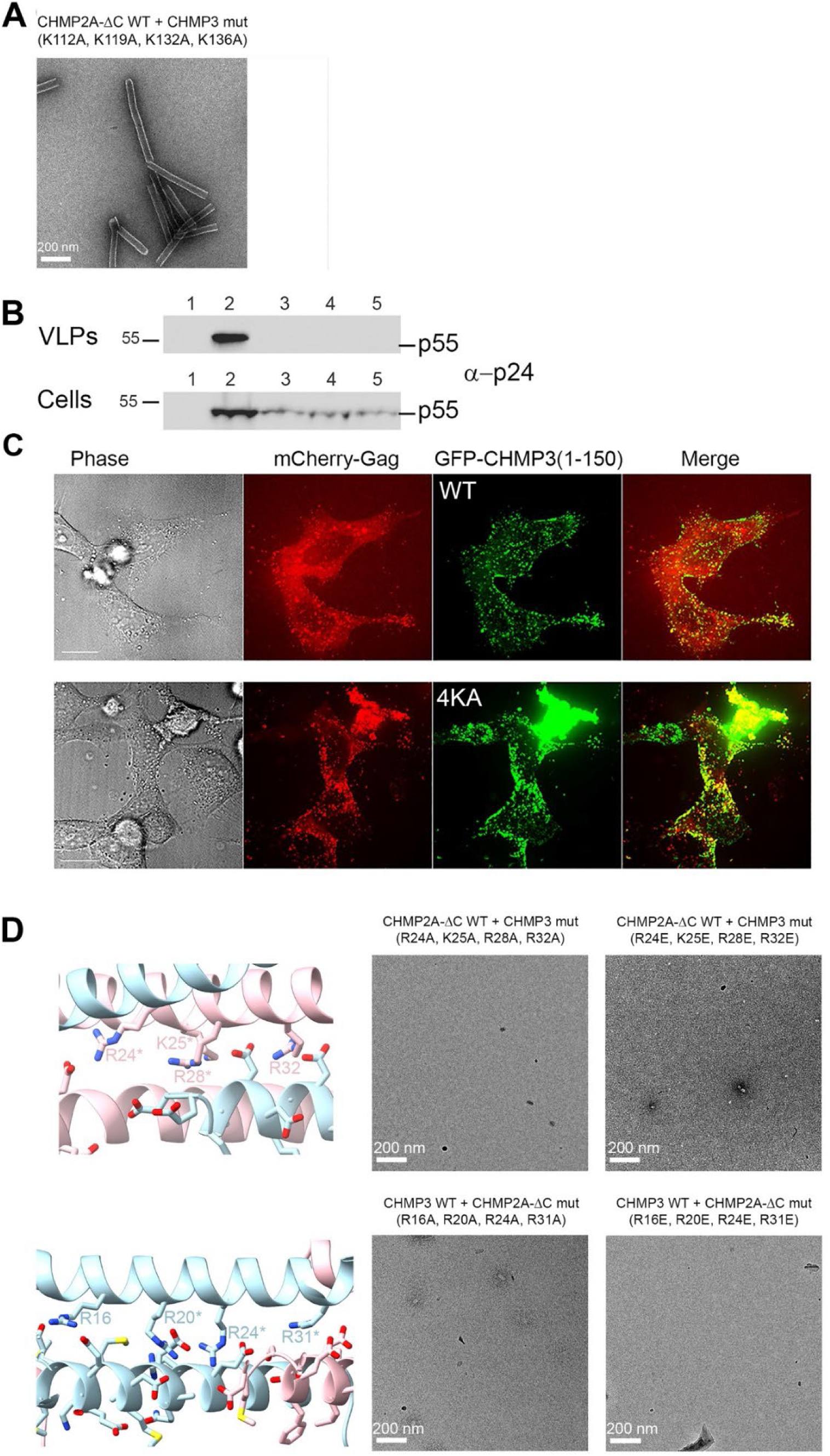
Structure-based mutagenesis of CHMP2A-CHMP3 polymer formation. **(A)** Negative staining electron micrograph showing regular tube formation by CHMP2A-CHMP3_K112A, K119A, K132A, K136A polymerization. Scale bar, 200 nm. **(B)** The substitution of four basic residues in GFP-CHMP3(1–150) 4KA (K112A, K119A, K132A, K136A) does not diminish its dominant-negative effect on HIV-1 budding. Western blot analyses of Gag released from Gag expressing cells as Gag-VLPs (upper panel) and detection of Gag in total cell extracts (lower panel): lane 1, Gag expression; lane 2, Gag and GFP-VPS4A E228Q expression; lane 3, Gag and GFP-CHMP3(1-150) expression; lane 4, Gag and GFP-CHMP3(1-150) 4KA expression. **(C)** Representative fluorescence images of HeLa cells transfected with Gag/mCherry-Gag and CHMP3(1-150) or CHMP3(1-150)4KA. Cellular distribution of wild-type and mutant KA GFP-CHMP3(1–150) indicates predominantly plasma membrane and intracellular localization as well as co-localization with mCherry-Gag. Scale bar, 10 µm. **(D)** Negative staining electron micrographs showing no polymer formation of CHMP3 mutants (upper left panel, close-up of a ribbon diagram illustrating the interface residues) R24A, K25A, R28A, R32A (upper middle panel) and R24E, K25E, R28E, R32E (upper right panel) with CHMP2A. (Lower panel) CHMP2A mutants (lower left panel, close-up of a ribbon diagram illustrating the interface residues) R16A, R20A, R24A, R31A (lower middle panel) and R16E, R20E, R24E, R31E (lower right panel) did not polymerize with CHMP3 *in vitro*. Scale bar, 200 nm.

**Figure S8:**
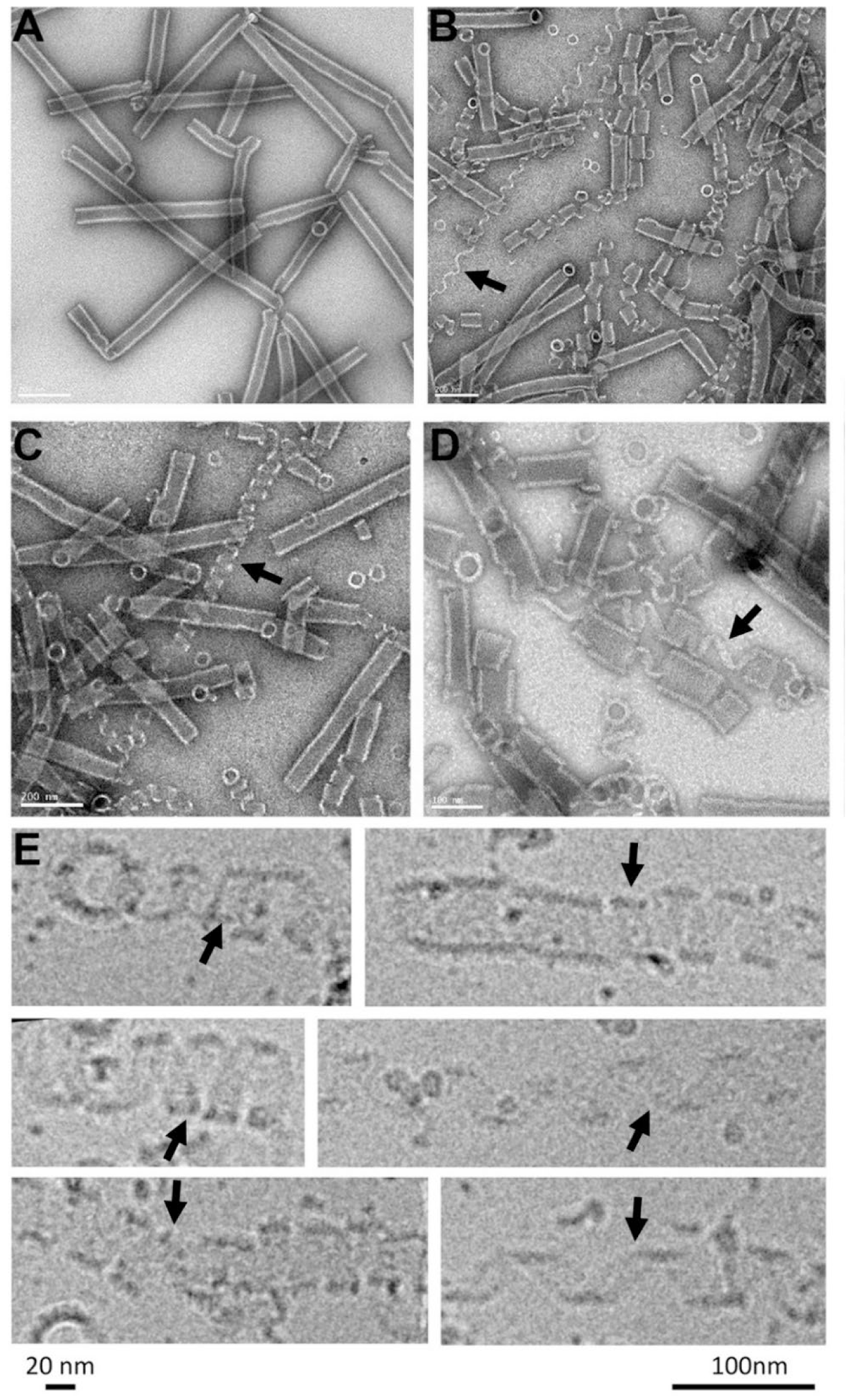
High ionic strength unwinds the CHMP2A-CHMP3 filaments. **(A)** Negative staining electron micrographs showing CHMP2A-CHMP3 wild-type polymers after treatment with 1M NaCl **(B, C)** and 1M KCl **(D)**. **(E)** Close-up of cryo-EM images shows unwinding of ∼20 nm wide filaments corresponding to the six-start helix observed in the structure. Single and multi-stranded unwound filaments are indicated by arrows.

**Figure S9:**
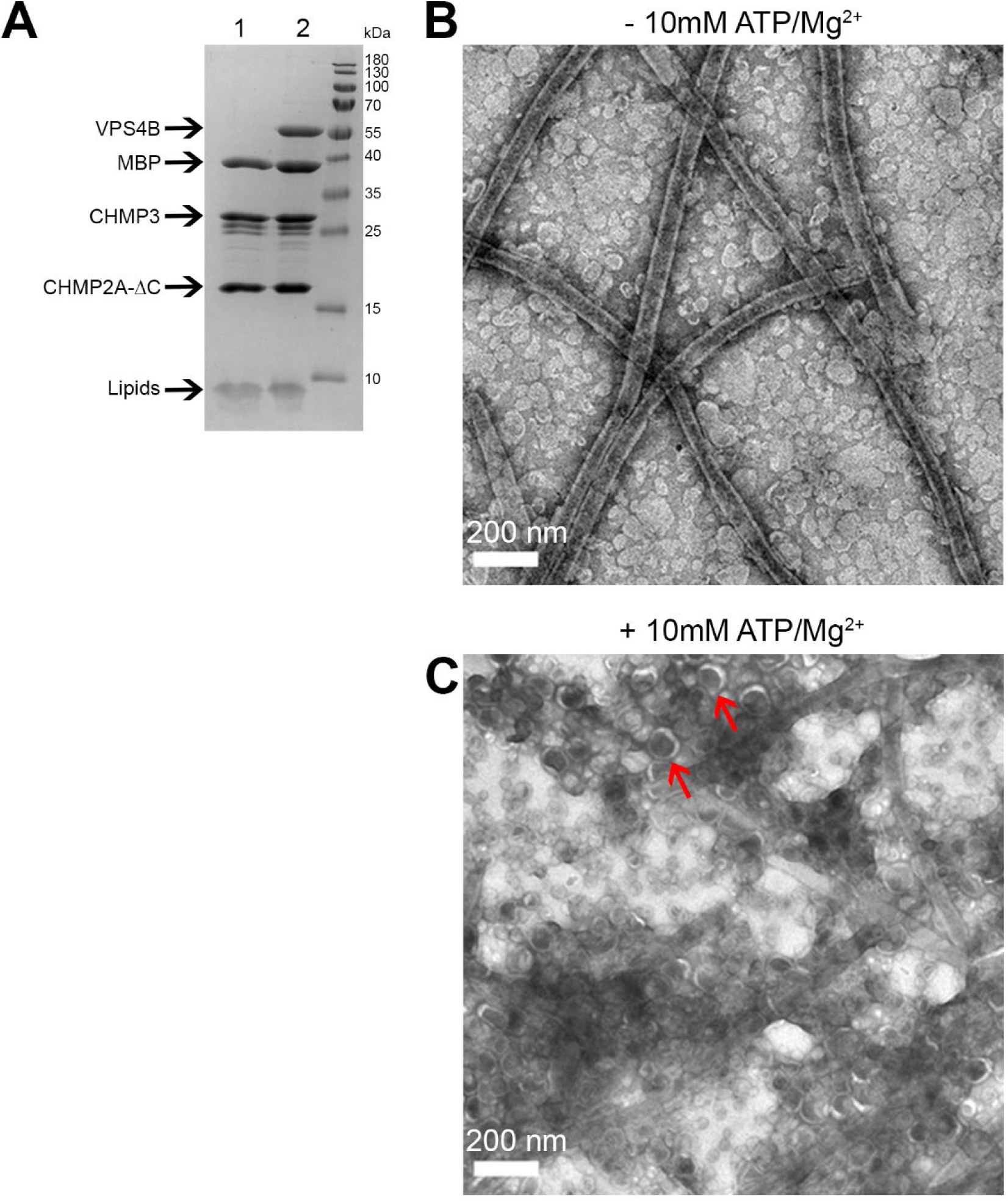
Incorporation of VPS4B and ATP into CHMP2A-CHMP3 membrane tubes induces their disassembly. **(A)** SDS-PAGE analyses of purified CHMP2A-CHMP3 polymers; lane 1, CHMP2A-CHMP3 polymers cleaved with TEV and coated with a lipid bilayer; lane 2 CHMP2A-CHMP3 polymers, TEV cleaved and incorporation of VPS4B prior to lipid bilayer coating. Negative staining electron micrographs of CHMP2A-CHMP3-VPS4B membrane-coated polymers before **(B)** and after **(C)** incubation with ATP and Mg^2+.^ Red arrows point to membrane vesicles resulting from tube cleavage. Scale bar, 200 nm.

**Figure S10:**
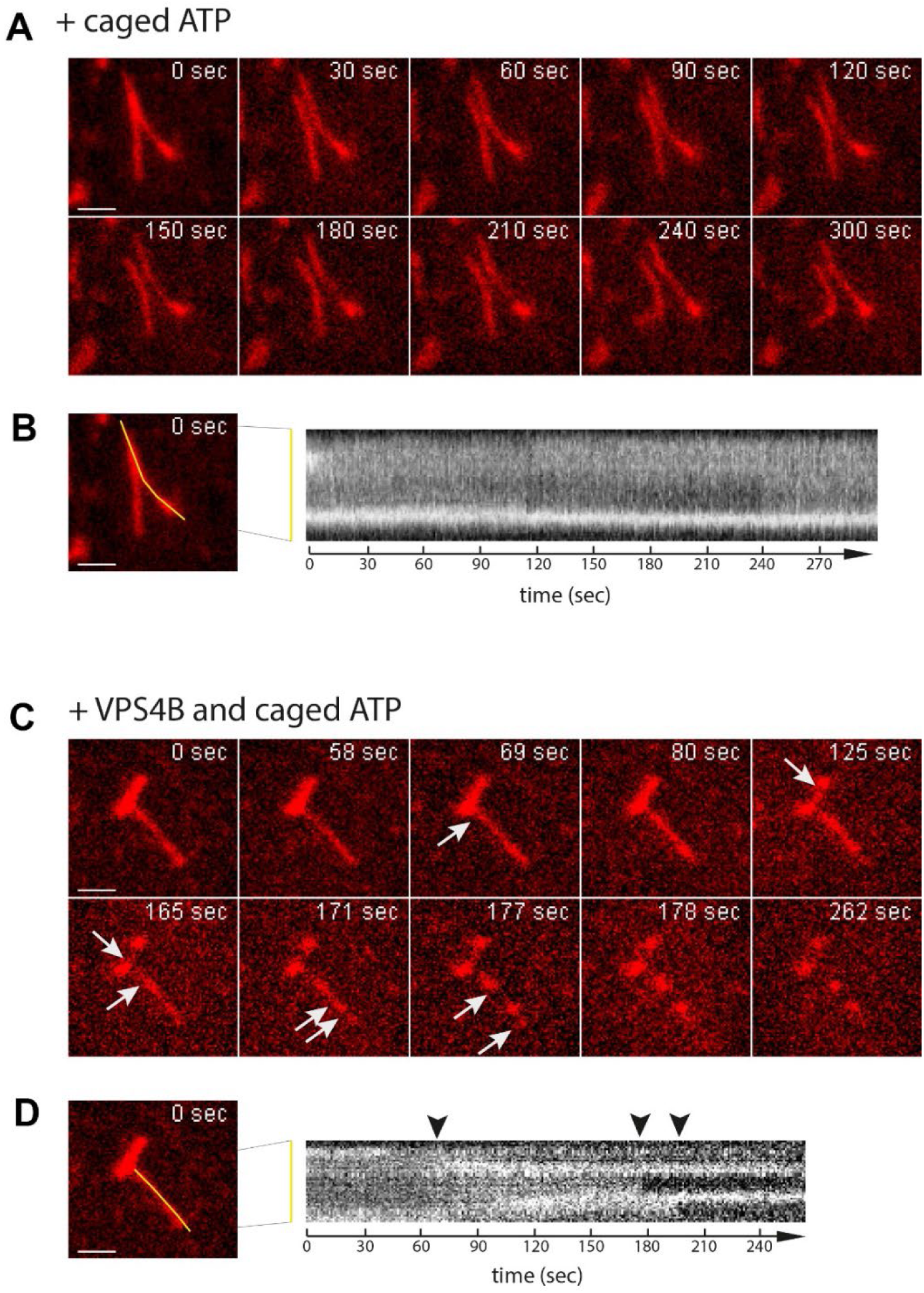
Imaging of VPS4B and ATP induced cleavage of CHMP2A-CHMP3 membrane coated tubes. **(A)** A CHMP2A-CHMP3-caged ATP membrane coated tube was activated at 365 nm (365 nm 30%, 100 ms at each time point) to uncage ATP and imaged over 282s, which indicated that uncaging did not change the tube structure (movie S3). **(B)** The kymograph of the tube (yellow line) shows that the tube stays intact over the imaging time. **(C)** Imaging of a CHMP2A-CHMP3-VPS4B-caged ATP membrane-coated tube following ATP uncaging (365 nm, 30%, 100 ms at each time point) reveals cleavage of the tube at several sites over the imaging time (movie S5). Scale Bar, 1µm. **(D)** The kymograph of the tube (yellow line) indicates tube cleavage (arrows) over the imaging time.

**Figure S11.**
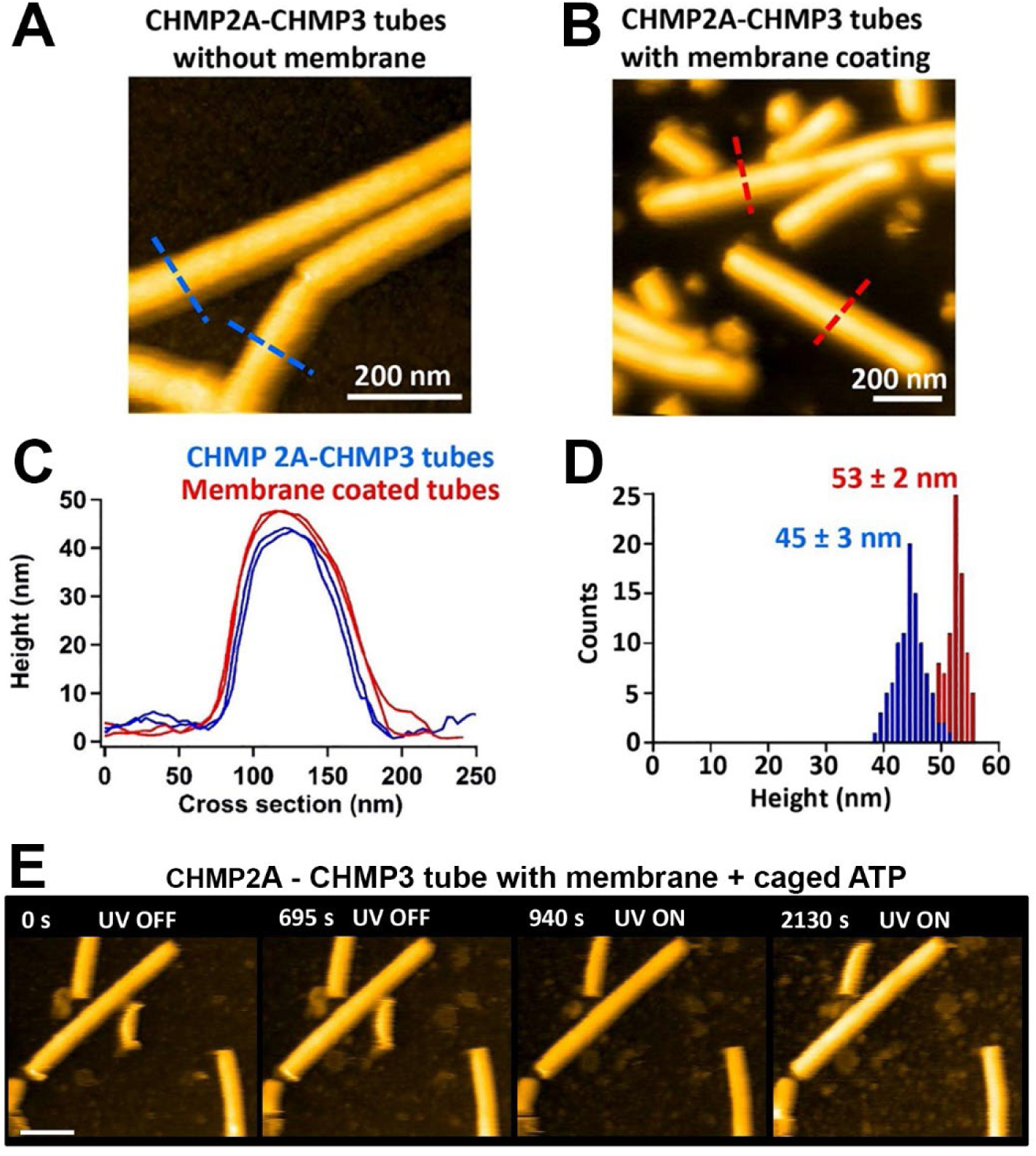
Height distribution of CHMP2A-CHMP3 tubes with and without membrane-coating. **(A)** AFM image of CHMP2A-CHMP3 tubes without membrane. **(B)** AFM image of CHMP2A-CHMP3 tubes coated with membrane. **(C)** Cross-section of AFM images of CHMP2A-CHMP3 in panel A (blue dotted line) and panel B (red dotted line). **(D)** Height histogram of CHMP2A-CHMP3 tubes with (red) and without (blue) membrane coating with respect to the surface. **(E)** Snapshots HS-AFM images of membrane-coated tubes loaded with 10 mM caged ATP taken with and without UV irradiation. The UV was turned on from 700 s onwards. Scale bar, 200 nm.

## Movies

**Movie 1: Fluorescence microscopy imaging of CHMP2A-CHMP3 membrane coated tubes.** A CHMP2A-CHMP3-caged ATP membrane-coated tube was activated at 365 nm for 10s to uncage ATP and imaged over 291s.

**Movie 2: Fluorescence microscopy imaging of CHMP2A-CHMP3 membrane coated tubes.** Another dataset of CHMP2A-CHMP3-caged ATP membrane-coated tube activated at 365 nm for 100 ms at each time point to uncage ATP and imaged over 300s.

**Movie 3: Fluorescence microscopy imaging of CHMP2A-CHMP3 membrane coated tubes.** A CHMP2A-CHMP3-VPS4B membrane-coated tube was activated at 365 nm for 10s to uncage ATP and imaged over 251s.

**Movie 4: Fluorescence microscopy imaging of CHMP2A-CHMP3 membrane coated tubes.** Imaging of a CHMP2A-CHMP3-VPS4B-caged ATP membrane-coated tube following ATP uncaging (365 nm, 10s). This demonstrates tube fission followed by a shrinking event from both sides

**Movie 5: Fluorescence microscopy imaging of CHMP2A-CHMP3 membrane coated tubes.** Another dataset of CHMP2A-CHMP3-VPS4B-caged ATP membrane-coated tube following ATP uncaging (365 nm, 100 ms at each time point) reveals cleavage of the tube at several sites over the imaging time.

**Movie 6: Fluorescence microscopy imaging of CHMP2A-CHMP3 membrane coated tubes.** Imaging of a CHMP2A-CHMP3-VPS4B-caged ATP membrane-coated tube following ATP uncaging (365 nm) showing at 36s a shrinking event from the end of a tube and at 51s a cleavage of the tube.

**Movie 7: HS-AFM imaging of CHMP2A-CHMP3 membrane-coated tubes.** HS-AFM movie of membrane-coated CHMP2A-CHMP3 tubes loaded with 10 mM caged ATP, taken before and after UV (365 nm) irradiation. Imaging time 5 seconds/frame.

**Movie 8: HS-AFM imaging of CHMP2A-CHMP3 membrane-coated tubes.** HS-AFM movie of membrane-coated CHMP2A-CHMP3 tubes loaded with 5 µM VPS4B and 10 mM caged ATP, taken without UV irradiation. Imaging time 3 seconds/frame.

**Movie 9: HS-AFM imaging of CHMP2A-CHMP3 membrane-coated tubes.** HS-AFM movie of membrane-coated CHMP2A-CHMP3 tubes loaded with 5 µM VPS4B and 10 mM caged ATP, taken with 365 nm UV irradiation. Imaging time 2 seconds/frame.

---

**Table S1.** Cryo-EM data collection, refinement and validation statistics.

|  | Membrane-bound CHMP2A<br>+ CHMP3<br>(Tube Diameter 430 Å)<br>(EMD-14630, PDB 7ZCG ) | Membrane-bound<br>CHMP2A + CHMP3<br>(Tube Diameter 410 Å)<br>(EMD-14631, PDB 7ZCH) |
| --- | --- | --- |
| **Data collection and processing** |  |  |
| Magnification | 130,000x | 130,000x |
| Voltage (kV) | 300 | 300 |
| Electron exposure per frame (e<sup>-</sup>/Å<sup>2</sup>) | 0.96 | 0.96 |
| Number of frames | 25 | 25 |
| Defocus range (μm) | 0.5-1.5 | 0.5-1.5 |
| Pixel size (Å) | 1.052 | 1.052 |
| Refined helical symmetry | 54.37°, 2.74 Å | 57.04°, 1.44 Å |
| Point group symmetry | C2 | C1 |
| Initial particle images (no.) | 45,847 | 80,524 |
| Final particle images (no.) | 25,353 | 11,396 |
| Map resolution (Å) | 3.3 | 3.6 |
| FSC threshold | 0.143 | 0.143 |
| **Refinement** |  |  |
| Initial model used (PDB code) | *de novo* | *de novo* |
| Model resolution (Å) | 3.6 | 4.0 |
| FSC threshold | 0.5 | 0.5 |
| Map sharpening *B* factor (Å<sup>2</sup>) | -96.57 | -101.52 |
| Model-to-map fit |  |  |
| Correlation coefficient, CC (mask) | 0.80 | 0.77 |
| Model Composition |  |  |
| Chains | 22 | 22 |
| Nonhydrogen atoms | 27,148 | 27,148 |
| Protein residues | 3,366 | 3,366 |
| *B* factor (Å<sup>2</sup>) |  |  |
| Protein | 74.18 | 90.17 |
| R.m.s. deviations |  |  |
| Bond lengths (Å) | 0.004 | 0.004 |
| Bond angles (°) | 0.758 | 0.762 |
| Validation |  |  |
| MolProbity score | 1.86 | 1.89 |
| Clashscore | 15.23 | 21.23 |
| Rotamer outliers (%) | 0.00 | 0.00 |
| Cbeta deviations (%) | 0.00 | 0.00 |
| Ramachandran plot |  |  |
| Favored (%) | 97.02 | 97.68 |
| Allowed (%) | 2.98 | 2.32 |
| Disallowed (%) | 0.00 | 0.00 |

